# Extracellular vesicles from a model of melanoma cancer-associated fibroblasts induce changes in brain microvascular cells consistent with pre-metastatic niche priming

**DOI:** 10.1101/2025.05.09.651827

**Authors:** M. Shelton, C. A. Anene, J. Nsengimana, M. K. Eldahshoury, B. Keane, W. Roberts, J. Newton-Bishop, J. R. Boyne

## Abstract

Malignant melanoma has one of the lowest 5-year survival rates of any cancer, and is recognised for being particularly invasive and metastatic, with the poorest survival outcomes in brain metastases patients. A key characteristic of these tumours is crosstalk between melanoma cells and cells of the tumour microenvironment (TME), such as cancer associated fibroblasts (CAFs). The role of melanoma-derived small extracellular vesicles (sEVs) in potentiating CAFs has been studied extensively, however the role of CAF sEVs in regulation of the local TME and distal pre-metastatic niche (PMN) is less clear. Here, we demonstrate that sEVs derived from an *in vitro* model of melanoma CAFs alter melanoma to promote oncogenic parameters within models of the TME and target a model of the brain PMN to promote changes associated with melanoma extravasation. Cargo profiling of these sEVs found significant differential expression of proteins, and RNA associated with pre-metastatic niche remodelling and unfavourable outcomes in patients. Together these data suggest a role for CAF sEVs in local and distal PMN formation, highlighting a potential therapeutic target for metastatic melanoma and identifying prospective liquid biomarker reservoirs.

## Introduction

Melanoma is the 5th most common cancer diagnosed in the UK, and incidence continues to rise year-on-year, with a projected increase of 9% in the UK between 2023-2025 [1]. Amongst solid tumours, melanoma has one of the highest propensities to metastasise to the brain with autopsy data indicating that around 75% of melanoma patients die with brain metastases [2]. Melanoma brain metastasis is predictive of a poor outcome despite advances in systemic therapy. This is reflected by brain metastases having the highest AJCC melanoma stage (Stage IV M1d). Patients with Stage IV melanoma and no brain metastases (Stage IV M1c or less) have an overall survival rate of 58% at 3 years with the most effective melanoma treatment, a combination immunotherapy regime of Ipilimumab and Nivolumab [3]. This compares to a 3-year overall survival rate of only 36.6% in patients with symptomatic brain metastases receiving the same treatment [4]. Not only do brain metastases predict the worst outcome for patients, but we lack biomarkers predicting metastasis to this site and response to immunotherapy.

Recent work has shed some light on the characteristics of melanoma brain tumours [5], but very little is known about the reasons for melanoma’s spread to the brain. Cancer-associated fibroblasts (CAFs) are key components of the melanoma tumour microenvironment (TME) and have been shown to drive disease progression, metastasis, and immunotherapy resistance [6–12]. CAFs exert these effects via their mechanical properties [13], secretion of cytokines, chemokines and other effector molecules [14], and also through the release of extracellular vesicles (EVs), a group of membranous particles that are released from almost all cell types and found in most biological fluids [15]. Small EVs (sEVs, also known as exosomes) have been the focus of significant research given their capacity to induce molecular changes via direct interaction with extracellular receptors on the cell’s surface, or become internalised and release biomolecular cargo into the intercellular space [16]. A series of elegant studies have demonstrated sEV-mediated regulation of the local TME in melanoma via modulation of melanoma metabolism [17], angiogenesis [18], and in a seminal piece of work, pro-oncogenic immunoregulation [19]. A key aspect of sEV-mediated cell-cell communication is that crosstalk can occur over much larger distances than is possible via bilayer-unprotected metabolites [20], facilitated by sEVs entering both the lymphatic and blood circulatory systems. Indeed, there is a growing body of evidence demonstrating that melanoma-derived sEVs are important players in the establishment of the distant pre-metastatic niche (PMN) [21]. Compared with cancer cell-derived sEVs, far less is understood about how CAF-derived sEVs influence the establishment of local and distant PMN in melanoma.

The aim of this study was to delineate the influence of CAF-derived sEVs on PMN formation and understand the underlying non-cell autonomous mechanisms of changes to cellular components of the melanoma TME in the brain. Specifically, we have established and validated an *in vitro* model of normal dermal fibroblast (NDF) re-education towards a CAF-like phenotype, and the successful isolation and characterisation of sEVs released by these cells. Here, we show that our model CAF sEVs are internalised and reprogram melanoma cells towards a more migratory and invasive phenotype while also driving pro-tumourigenic changes in other cell types found in the local TME. Moreover, we show that CAF sEVs promote remodelling of hCMEC/D3 brain endothelial cells, an early event associated with establishment of the brain PMN. To understand the role of CAF sEV cargo in these phenotypic changes, we performed small RNA-seq and quantitative proteomic profiling, revealing a signature of sEV miRNA and proteins that include reported drivers of endothelial to mesenchymal transition (EndMT). Taken together, our data show that reprogrammed CAFs secrete sEVs that promote metastasis, remodel brain endothelial cells, and carry a distinct biomolecular signature linked to endothelial-mesenchymal transition.

## Materials and Methods

### Cell culture

NDFs were prepared from skin specimens collected from the Huddersfield Royal Infirmary through routine abdominal surgical procedures with National Health Service (NHS) Research Ethics Committee (REC) approval (ref no: 15/EM/0265) and informed consent from patients with no history of skin malignancy and isolated as described [22]. Human melanoma cell lines A2058 and A375 were purchased from European Collection of Authenticated Cell Cultures. Primary HUVECs were purchased from Thermofisher. The hCMEC/D3 cell line was a gift from Dr Paul Meakin (University of Leeds). All cell lines were routinely tested and certified mycoplasma free. Fibroblast and A2058 cells were maintained in High Glucose DMEM (Sigma Aldrich) and A375 cells in RPMI 1640 Medium with L-Glutamine (BioWhittaker), supplemented with 10% FBS (Sigma Aldrich) and 100U/mL penicillin/streptomycin (Gibco). Endothelial cells were grown in Endothelial Cell Growth Medium with SupplementMix (Promocell). Cells were routinely cultured at 5% CO_2_ and 37°C in T75 flasks (primary cell lines never grown to more than passage 5). Cells were counted using the Corning^®^ Automated Cell Counter (Corning). For generation of model CAFs, NDFs were seeded in flasks in 10% DMEM and left to attach and grow for at least 24 hours. Media was then changed to DMEM supplemented with 5ng/mL of recombinant human TGF beta 1 (TGFβ1) protein (Abcam) for 48h. For treatment of cells with sEVs, cells were grown (0.3-3 x 10^5^ cells) in 6 or 24-well plates for 24h in EV-free media, then treated with 3 x 10^3^ sEVs per cell, isolated from either A2058, A375, NDF or CAFs and incubated for 24-72h.

### sEV isolation

Cell lines were grown to ∼80% confluency, then grown in sEV free media for 16h. 40-80mL media was spun at 300x g then 4000x g and 0.2µm filtered, then concentrated to 0.5mL using the Amicon^®^ Ultra-15 10kDa Centrifugal Filter Unit (Sigma Aldrich) according to the manufacturer’s protocol. 0.5mL media was then run on a SEC qEVoriginal (Izon) according to manufacturer’s protocol, and fractions containing sEVs (fractions 7-11) were collected.

### Nanoparticle Tracking Analysis

NTA measurements were performed on a Malvern NanoSight NS300 (NanoSight, Amesbury) with a Blue488 laser type and sCMOS camera. NTA 3.2 Dev Build 3.2.16 software was used to capture and analyse the videos. All samples were diluted 1:20-100 in EV-free PBS (Gibco) and three measurements of 90 sec were carried out for each sample at 22°C with syringe speed 75.

### CellMask™ Orange sEV staining

5mg/mL of CellMask™ Orange Plasma membrane Stain was diluted 1:1000 in EV-free PBS (Gibco). 1:1 (v/v) of working stain was added to sEVs and incubated at 37°C for 10 min. sEVs underwent additional centrifugation and PBS wash steps with Amicon^®^ Ultra-0.5 100kDa Centrifugal Filter Unit (Sigma Aldrich) as described [23] to remove unbound dye. sEVs were quantified via NTA and an equal number of sEVs were added to cells and left to incubate for 24h. An equal volume of PBS with or without CellMask™ Orange was used as controls. Cells were then used for downstream analysis via flow cytometry or immunofluorescence microscopy.

### Transwell co-culture of cell lines

NDFs/CAFs were treated with 20μM of GW4869 (Sigma) for 24h prior to coculture, with sEV secretion depletion being confirmed via NTA. 5 x 10^4^ endothelial cells were seeded into the bottom of a 24 well plate. 5 x 10^4^ NDF/CAFs were seeded into 0.4µm transwell inserts (Sarstedt) and incubated in 24 well plates for 24h before lysate and RNA was isolated.

### Angiogenesis tube formation assay

96 well plates were coated with 30µL of Geltrex™ LDEV-Free Reduced Growth Factor Basement Membrane Matrix (Gibco) and incubated at 37°C for 30 min. Endothelial cells were seeded at 1.5 x 10^4^ cells per well. 25ng/mL VEGF-A165 (ThermoFisher) or 4% DMSO (ThermoFisher) were used as positive and negative controls of angiogenesis, respectively. Cells were left for 6-18h to form networks of branching structures, captured on the EVOS XL Core Cell Imaging System (10x objective). Images were quantitatively analysed using AngioTool.

### Transwell assay

Cells were serum starved for 24h prior to the assay. 1mL serum containing media was added to the lower chambers of a 24 well plate with 8µm TC-inserts (Sarstedt), and 500µL cell suspension (1 x 10^4^ cells) was added into the insert. Prior to seeding, inserts were coated with 50µL Geltrex™ LDEV-Free Reduced Growth Factor Basement Membrane Matrix (Gibco) and incubated at 37°C for 1h. Cells were incubated for 6-24h at 37°C, then fixed with ice cold 70% ethanol for 30 min. Insert membranes were stained in 0.2% crystal violet (Fisher Science) for 10 min, then mounted onto slides. 5 random high-power images were taken (EVOS XL Core Cell Imaging System, 4x objective) and an average was calculated using ImageJ.

### Collagen contraction assay

Fibroblasts were resuspended so 3 x 10^5^ cells were present in 1.2mL media. 600µL of 3mg/mL rat tail collagen I (ThermoFisher) was added to cells, then 21µL 1M NaOH. 500µL was added per well of a 24 well plate and allowed to solidify for 20 min at 37°C, then 500µL media was added to each well. Gels were dissociated carefully from well walls using a 10µL pipette tip and cells were left to contract for 24h. Images were quantified on ImageJ normalising area of gel outline to the area of the entire well.

### Mass Spectrometry

Protein from 500μL pooled sEVs from multiple NDF donors and their TGFβ1 transformed CAFs in duplicate were extracted using RIPA, as described in Supplementary Methods. Samples were analysed by Dr. Kate Heesom (University of Bristol) using tandem mass tagging coupled with liquid chromatography mass spectrometry (TMT LC-MS). The TMT-labelled proteomic dataset was processed to ensure the accuracy and reliability of protein identifications. Protein codes were cross-referenced with the UniProt database (www.uniprot.org) to verify their presence in the reviewed (Swiss-Prot) dataset, ensuring high-confidence annotations. Only proteins with valid UniProt accessions were retained for downstream analysis. The EXOCARTA dataset was downloaded from exocarta.org. Gene identifiers from Exocarta were converted to UniProt IDs using the UniProt ID mapping tool to standardise protein identifiers and facilitate comparison with the experimental dataset. The processed TMT proteomic dataset was compared against the EXOCARTA list to identify overlapping proteins. Gene ontology (GO) enrichment analysis was performed using the PANTHER classification system (version 19.0, Gene Ontology Consortium). Enrichment analysis was conducted against the Gene Ontology Biological Process (GO-BP) database using Fisher’s exact test with false discovery rate (FDR) correction to account for multiple testing. GO terms with an FDR-adjusted p-value <0.05 were considered significantly enriched. Search Tool for the Retrieval of Interacting Genes/Proteins (STRING) network analysis was used to generate predicted protein interactions. The mass spectrometry proteomics data have been deposited to the ProteomeXchange Consortium via the PRIDE partner repository [23] with the dataset identifier PXD063635.

### RNA sequencing

NDF/CAF sEVs were pooled from 6 donors in duplicate. 200ng of small RNA sample was sent to Novogene for SE50bp sequencing (raw data is available via GEO). Raw reads were subjected to additional adaptor trimming and low-quality reads were removed using Trimmomatic v0.39 with parameters (WindowSize = 4, Required Quality > 20). Trimmed reads were aligned to the GRCh38/hg38 assembly of the human genome using Bowtie v1.3.1 with parameters (-k 10 -v 0). Htseq-count v0.11.1 was used to quantify the miRNA abundance based on the human miRNA annotation (miRbase Release 22.1) with parameters (-s no -a 10 -m union --nonunique none). The expression levels of miRNA and mRNA were normalised by counts per million (CPM). Differential expression analyses between different groups were performed using the Limma-Voom method based on the Limma R package. For sEV samples, the differential expression was defined at p≤0.01 due to the small number of detected RNAs across the samples.

### Statistical analysis

Data are expressed as the mean ± standard error of the mean from at least two independent experiments carried out in triplicate. Statistical analysis was performed using GraphPad Prism 8.4.2. Principle component analysis and volcano plots were created using R 4.2.2. An unpaired t-test or one sample Wilcoxon Signed Rank test was used to compare the means of two independent groups. A one-way ANOVA or Kruskal-Wallis ANOVA was used to determine statistical significance between the means of three or more independent experimental groups, followed by a Tukey’s post-hoc test or unpaired t with Welch’s correction. Significant results are denoted as: p ≤ 0.05 = *, p ≤ 0.01 = **, p ≤ 0.001 = ***, p ≤ 0.0001 = ****, or p > 0.05 = ns (not significant).

## Results

### Establishment of a melanoma CAF model

CAFs are defined as persistently activated fibroblasts in the tumour stroma that influence many parameters of melanoma progression [24]. When NDFs transition into CAFs, they take on characteristics similar to smooth muscle, becoming more contractile, migratory and increasing secretion of various ECM remodelling proteins [25]. Whilst CAF populations between individuals and within the primary TME are heterogenous, expression of certain markers have been identified to signify CAF activation from NDFs [26]. CAF activation is a complex process that can occur via multiple routes and through physical and biochemical stimulatory factors, however, transforming growth factor β (TGFβ) is widely reported to act as a potent inducer of CAF differentiation [27–30]. To mitigate heterogeneity between patient samples and considering that CAFs are generally activated due to tumour-derived signalling rather than intrinsic mutations, we sought to generate an *in vitro* model that exhibited CAF-associated phenotypic properties and displayed characteristic markers.

To this end, primary NDFs were isolated from skin and cultured with TGFβ1 to re-educate cells to a CAF-like state. Consistent with previous observations [31, 32], TGFβ1 induced an increase in protein expression of α-SMA in treated NDFs, as evidenced by Western blot, IF and FACs analysis (Fig. 1A-C, Supplementary Fig. 1D). To determine if our model CAFs displayed reported transcriptional changes, we analysed the expression of four genes that are controlled by the TGFβ1 signalling cascade and upregulated in melanoma CAFs [31, 33]. As can be seen in Fig. 1D, we observed a significant increase in the transcript levels for each of these genes in our TGFβ1-treated model CAFs compared with untreated control. In addition, we also assessed the expression of several miRNAs previously reported to be differentially expressed in melanoma CAFs [34, 35]. In each case, we observed a significant difference in the abundance of these transcripts in our model CAF cells compared with control (Supplementary Fig. 1E).

**Figure 1:**
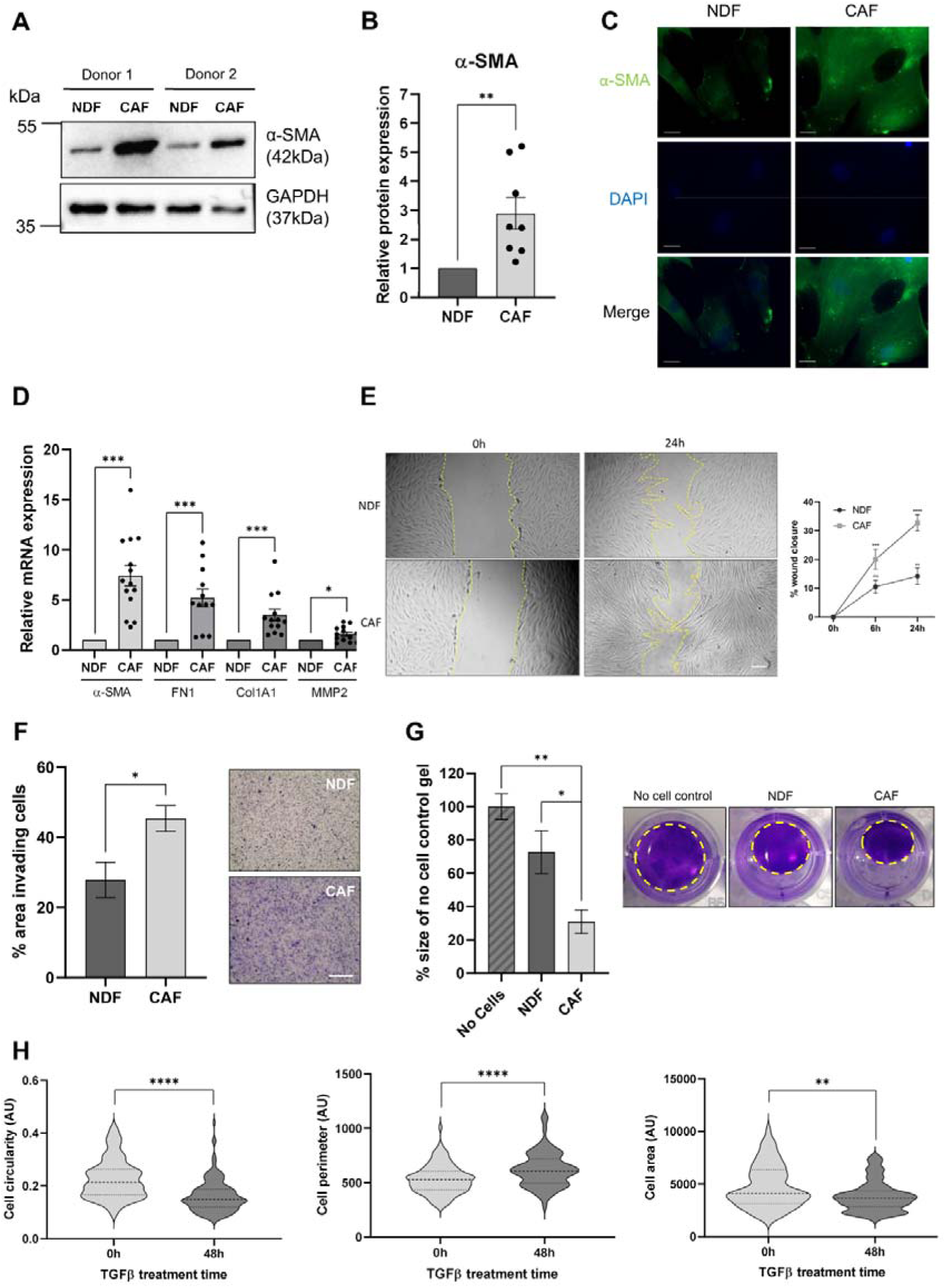
Characterisation of model CAFs treated with TGFβ1. **(A)** Representative western blot of α-SMA expression in two donor populations treated with or without 5ng/mL TGFβ1 for 48 hours. **(B)** Quantification of α-SMA protein expression normalised against GAPDH. **(C)** Representative immunofluorescence microscopy images of NDF or model CAFs stained with α-SMA-FITC and DAPI. Scale bar = 20µm. **(D)** Relative transcript expression of characteristic CAF markers normalised against reference genes (GAPDH, TBP and RER1) in NDFs compared with model CAFs. **(E)** Representative images of a scratch wound assay, % wound closure in NDFs and model CAFs. **(F)** Quantified cell invasion as % area of transwell containing invading cells per field of view and representative images of transwell invasion assay for NDFs and model CAFs. Scale bar = 500µm. **(G)** Quantification of collagen contraction over 24h, relative to no cell control and representative images of collagen gel contraction of NDFs or model CAFs. Yellow dashed line = outline of gel. **(H)** Cell circularity, cell perimeter and cell area quantified in ImageJ at 0h and 48h post TGFβ1 treatment. Dashed lines show median and quartiles.

Next we investigated changes in several phenotypic parameters consistent with reported *in vivo* characteristics of CAFs, including increased cell migration, invasion and crucially contractility within a 3D-matrix [36]. As can be seen in Fig. 1E-G, we observed a significant shift towards CAF-like properties. Moreover, model CAFs had significant increases in ATP production and mitochondrial dehydrogenase activity, despite no changes to cell viability, suggesting increased proliferation (Supplementary Fig. 1A-C). Finally, morphological analysis of model CAFs reflected myofibroblast-like changes, with longer protrusions giving decreased circularity and increased perimeter (Fig. 1H). Together, these data demonstrate that TGFβ1-treatment of primary NDFs generate CAF-like cells that display biochemical and phenotypic parameters consistent with melanoma CAFs.

### Model CAFs release sEVs that are internalised by cells typical of the local and distal melanoma TME

We next sought to isolate and characterise sEVs released by our model CAF cells, observing MISEV guidelines [37]. To this end, TGFβ-induced model CAF cells were cultured in EV-free media prior to isolation of sEVs via size-exclusion chromatography (SEC), which facilitates a high yield of sEVs, low variability between isolations, and a homogenous population with retained functionality [38]. As seen in Fig. 2A, particles were enriched in SEC fractions 7-11, and downstream analysis using NTA and TEM showed that particles from these collated fractions displayed an average size that fell within the 30-150nm range characteristic of sEVs (Fig. 2B + Supplementary Fig. 2A+B). To confirm that isolated particles harboured reported sEV marker proteins, CD63 and CD81, western blot analysis was performed. As can be seen in Fig. 2C particles from SEC fractions 7-11 were enriched in sEV markers while in contrast the negative sEV marker, Calnexin, was absent, demonstrating that isolated particles were not contaminated with membranous structures derived from the intracellular endoplasmic reticulum. We next sought to confirm the presence of the canonical sEV marker protein, CD63, at single particle level via imaging flow cytometry and in line with MIFlowCyt-EV guidelines [39]. As can be seen in Fig. 2D, we observed a robust signal for CD63 on single particles and when plotting the intensity of gated particles, there is a clear population of CD63^+^ particles within the total population (Supplementary Fig. 2C+D).

**Figure 2:**
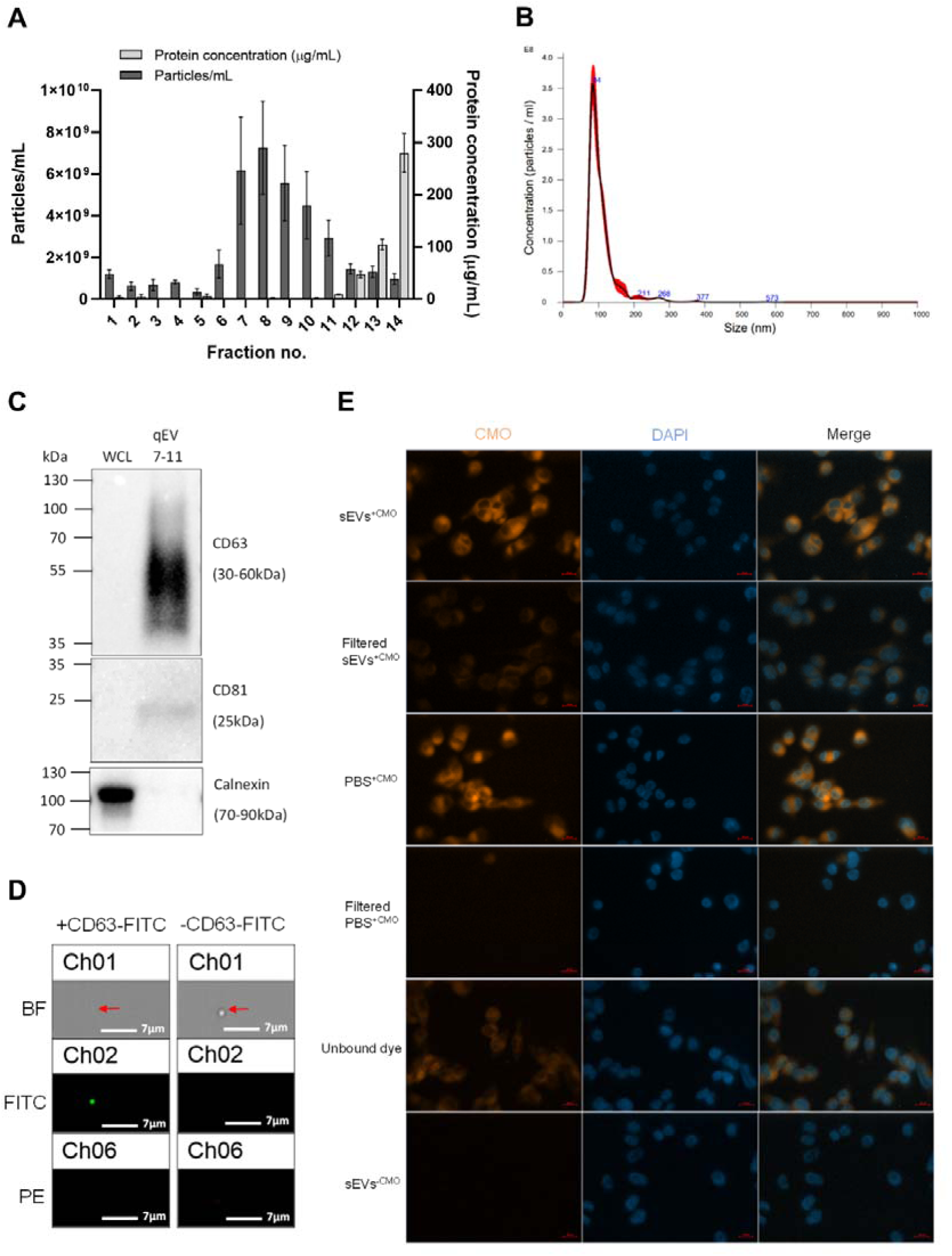
Characterisation of CAF sEVs isolated via SEC and their uptake into recipient cells. **(A)** Average particle and protein concentration of fractions 1-14 eluted from SEC of model CAF cell cultured media measured via NTA and micro-BCA, respectively. Three 90 second videos were captured per sample replicate for NTA. **(B)** A representative NTA size and concentration profile of combined fractions 7-11 from model CAFs isolated via SEC. Red error bars indicate SEM. **(C)** Western blot of model CAF SEC combined fractions 7-11 alongside model CAF whole cell lysate (WCL) against sEV markers. **(D)** Representative FlowSight images of sEVs stained with or without CD63-FITC after gating for EVs based on area and aspect ratio. Scale bars = 7µm. **(E)** Representative IF microscopy images of A375 cells co-cultured with model CAF sEVs or EV-free PBS stained with or without CMO, with or without unbound dye removed. Scale bar = 20µm.

In order to investigate if model CAF-derived sEVs are internalised when co-incubated with target cells, sEVs were stained with amphipathic CellMask™ Orange (CMO) dye [40] and CMO binding to sEVs confirmed via amplified fluorescence polarisation compared with the unbound dye (Supplementary Fig. 2E). Stained sEVs were then co-cultured with a range of cell lines that are representative of the melanoma TME, including the melanoma cell lines A375 and A2058, NDFs, HUVECs (as a model of local TME endothelial cells) and hCMEC/D3 (as a model of brain endothelial cells). In each case, fluorescence microscopy imaging showed diffuse CMO staining within cells co-incubated with labelled sEVs, with a higher fluorescence signal in filtered sEVs compared to filtered PBS (Fig. 2E + Supplementary Fig. 2F), data that was also confirmed and quantified via FACS analysis (Supplementary Fig. 2G-L). Together these data demonstrate that model CAFs release sEVs that are internalised by a range of cells commonly associated with the melanoma TME.

### Model CAF sEVs drive re-education of melanoma TME cells *in vitro*

After confirming that melanoma and other local TME cells internalise model CAF-derived sEVs, we investigated how this uptake affects recipient cell properties compared to the uptake of sEVs from NDFs. In addition, as previous studies have shown that melanoma-derived sEVs play a role in re-educating cells within the TME [41] we also decided to isolate sEVs from the A375 and A2058 melanoma cell lines, which exhibit low and high invasive potential, respectively [42, 43] and determine how these might impact the NDF-CAF axis. Initially, sEVs derived from each of these cell backgrounds were added to NDFs to determine how they impacted on fibroblast activation. As can be seen in Fig. 3A-E, sEVs from CAFs and the invasive melanoma line A2058 significantly increased proliferation, migration, and contraction of NDFs, when compared with sEVs isolated from NDFs and A375 cells. Moreover, CAF derived sEVs caused significant increases in transcript levels of α-SMA and FN1 (Fig. 3F) but did not significantly increase levels of MMP2 or Col1A1 (Supplementary Fig. 3C). Next, we examined how co-culture with sEVs impact the less invasive A375 melanoma cell line. Interestingly, we observed that model CAF-derived sEVs caused a significant increase in the invasion potential of A375 cells (Fig. 3G), but not migration in the absence of basement membrane (Supplementary 3D). Analysis of EMT transcript levels revealed increased levels of vimentin transcripts in cells co-incubated with CAF-derived sEVs (Fig. 3H), however, there were no significant changes to SNAIL or TWIST (Supplementary Fig. 3E). Taken together, these studies suggest that CAF-derived sEVs can alter the metastatic properties of melanoma cells and re-educate NDFs towards a CAF-like phenotype, indicating a potential multi-directional crosstalk facilitated by sEVs released from both melanoma and CAF populations within the TME.

**Figure 3:**
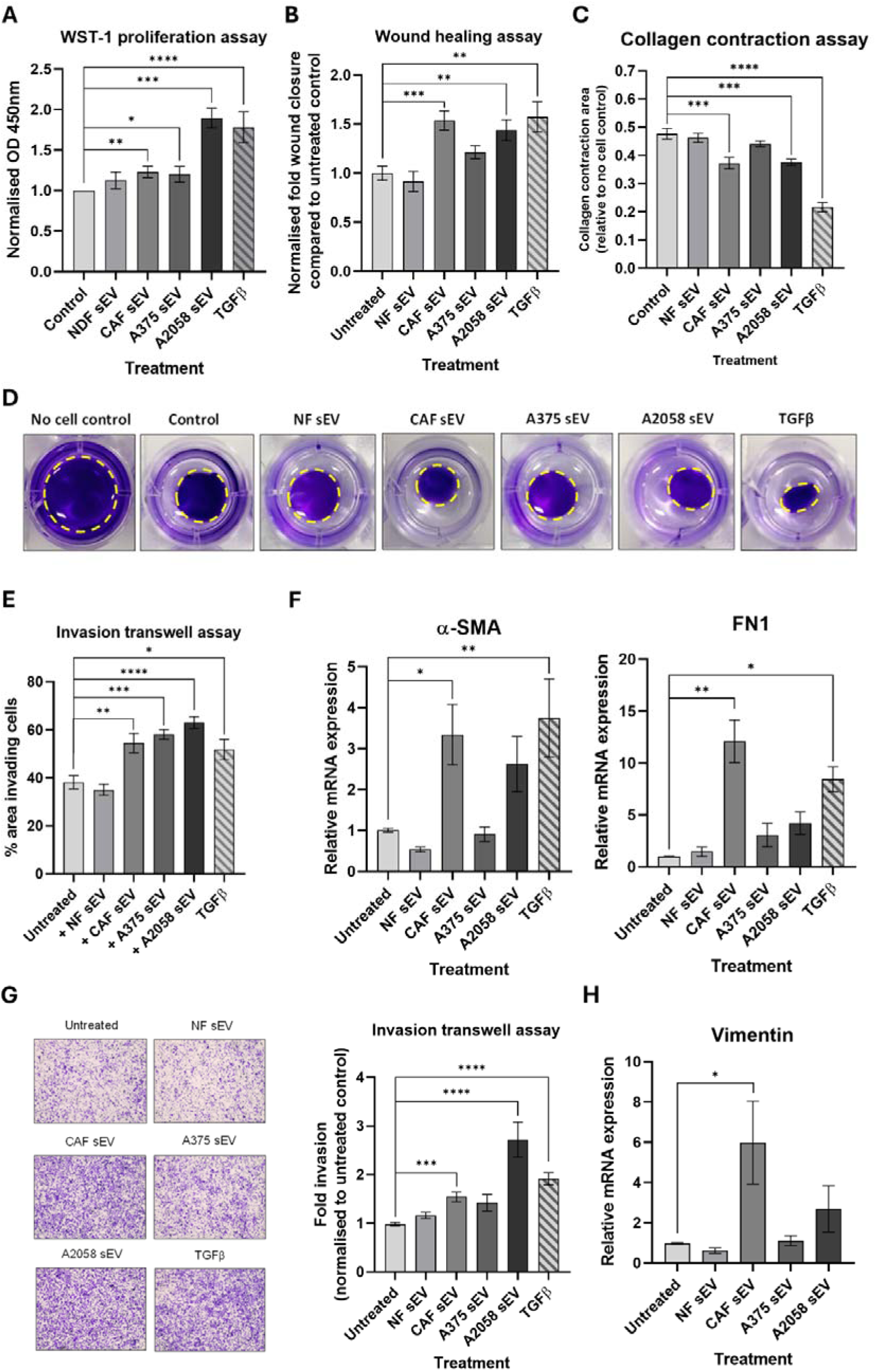
CAF sEV treatment affects multiple phenotypic parameters in dermal fibroblasts and melanoma cell line A375. **(A)** NDF co-cultured with sEV were analysed via WST-1 assay, and normalised fibroblast proliferation rate determined as optical density at 450nm using 630nm as a reference to measure formazan dye production. **(B)** Quantified sEV-treated NDF migration compared with untreated control after 24h. **(C)** Collagen matrix contraction after 24h co-culture of NDFs with sEVs relative to no cell control. **(D)** Representative images of NDF collagen contraction assay 24h post indicated treatment (yellow dashed line = outline of gel). **(E)** Quantified cell invasion as fold invasion through transwell per field of view normalised to untreated NDF control. **(F)** Relative transcript levels of CAF markers α-SMA and FN1 in sEV treated NDFs. **(G)** Representative 24h images of A375 transwell invasion assay alongside quantified cell invasion as fold invasion through transwell per field of view normalised to untreated A375 control. **(H)** Relative expression of vimentin in sEV-treated A375 cells.

### Model CAF sEVs drive an angiogenic switch in endothelial cells

Given our *in vitro* data demonstrating sEV multi-directional crosstalk between melanoma cell and model CAFs, we next sought to elucidate whether melanoma and CAF sEVs might also impact on a model of local TME endothelia. To this end, HUVEC cells were cultured with sEVs, and a tube formation assay performed to assess angiogenic stimulation. As can be seen in Fig 4A-C, we observed a significant increase in branching index (indicative of a potent pro-angiogenic stimuli) when HUVECs were co-incubated with either model CAF or A2058-derived sEVs, that was absent for NDF-derived sEVs. Next, we assessed the impact of CAF sEV co-incubation on the abundance of a pro-angiogenic proteins in HUVECs (Fig 4D-F). We observed increased levels of numerous pro-angiogenic proteins including TIMP-1, a factor upregulated in colorectal cancer EVs that induces ECM remodelling in recipient fibroblasts [28]. In the context of cancer progression endothelial to mesenchymal transition (EndMT) is a key process in endothelial remodelling that compromises vascular integrity and promotes mesenchymal traits, enabling increased permeability and matrix remodelling [44]. To investigate if CAF-derived sEVs were promoting EndMT, mesenchymal markers were assessed via qRT-PCR (Fig. 4G-K + Supplementary Fig. E-G). We observed a significant increase in α-SMA transcript abundance in HUVECs treated with model CAF sEVs, but no other EndMT markers reached significance. Together, these results demonstrate that model CAF sEVs induce angiogenic potential and expression of pro-angiogenic and EndMT markers in HUVECs.

**Figure 4:**
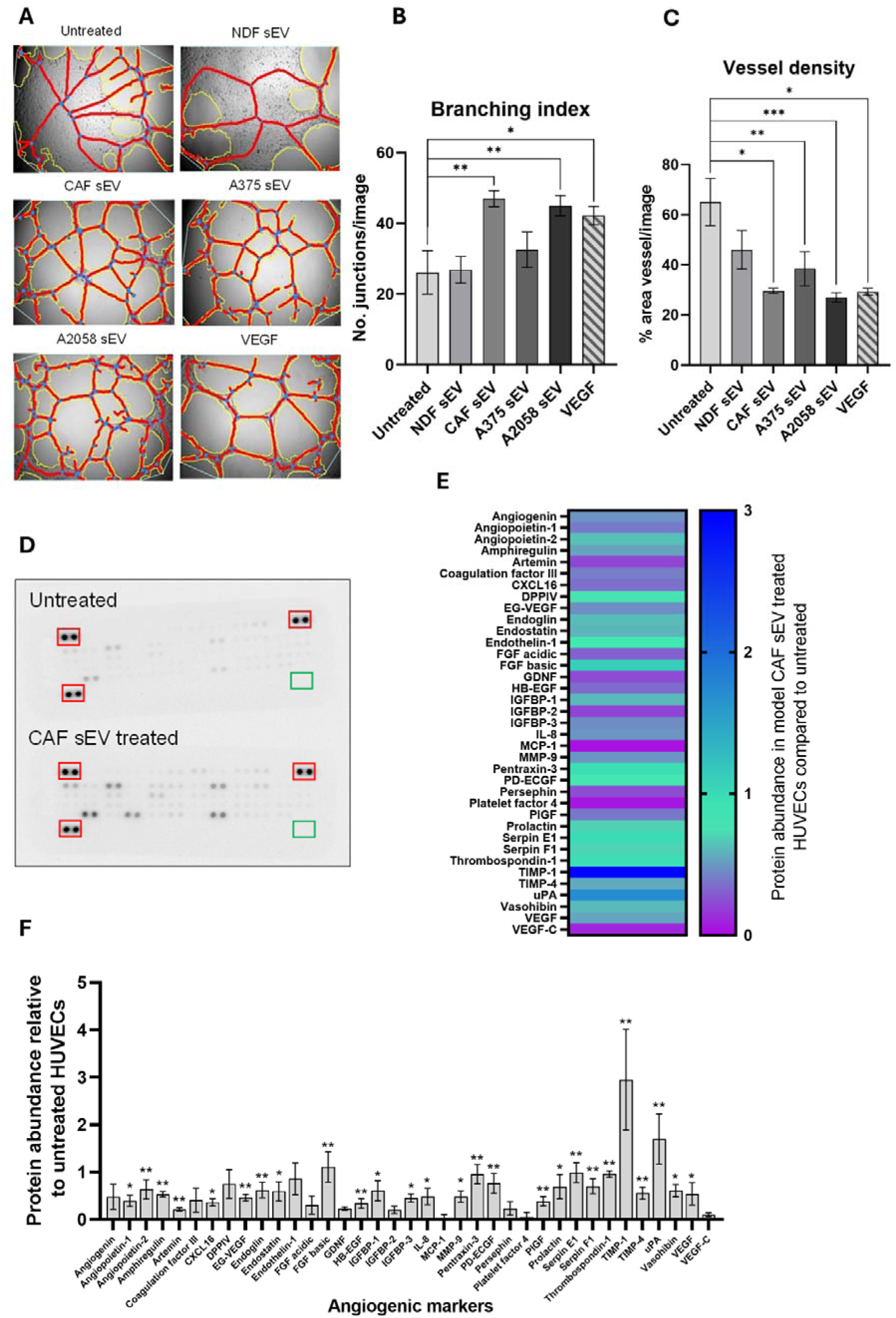
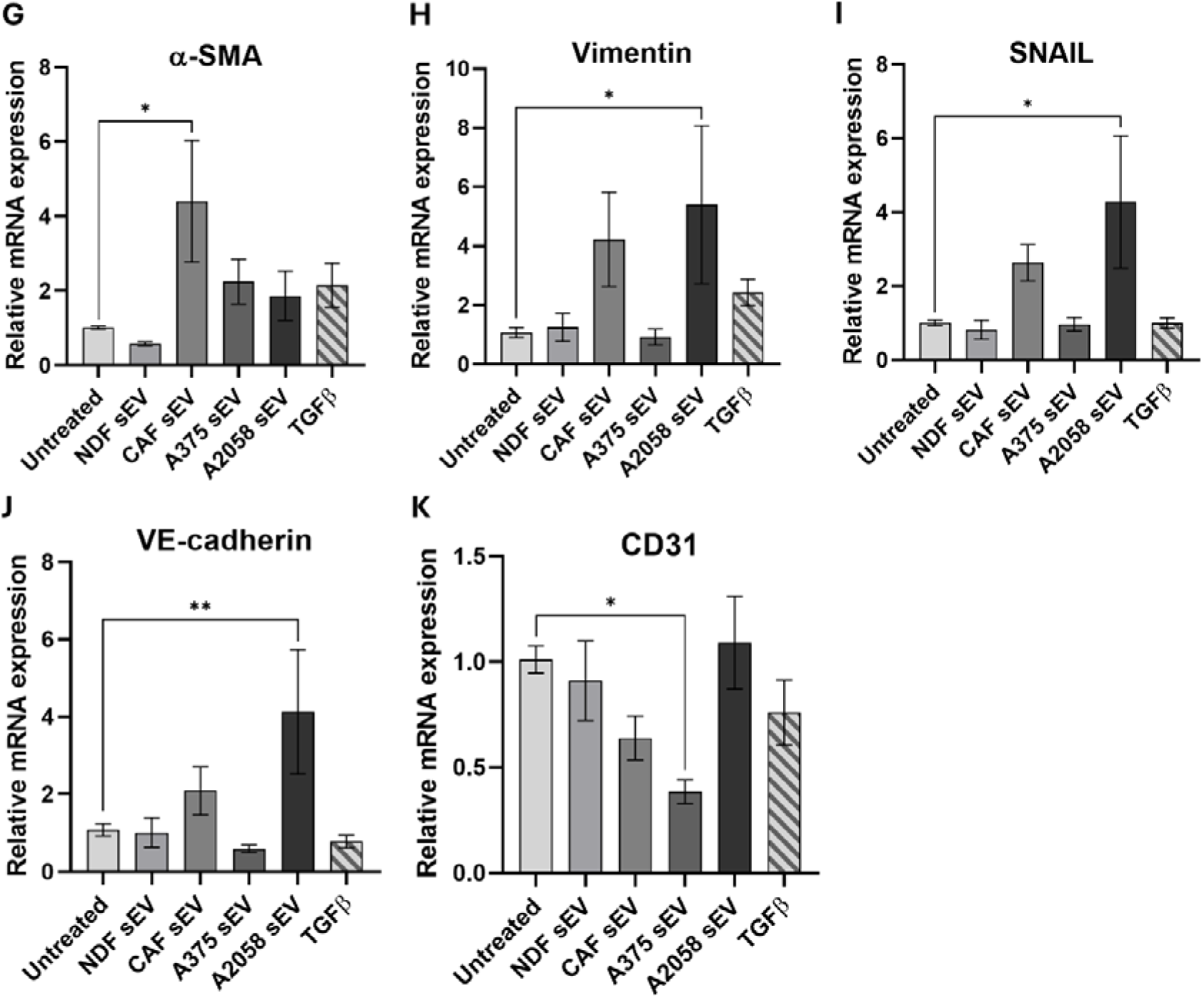
Model CAF sEV treatment changes angiogenic properties of HUVECs. **(A)** Representative angiogenesis branching assay images for HUVECs with treatment of sEVs or VEGF. Images were captured on the EVOS XL Core Cell Imaging System, at 10x objective and analysed using AngioTool (white line = explant area, yellow line = boundary of vessel, red line = vessel, blue dot = branching point). Quantified **(B)** branching index (no. of branching junctions per image) and **(C)** Vessel density (% area in explant area occupied by vessels). Statistical significance was determined for all using a one-way ANOVA or a Kruskal-Wallis ANOVA, respectively. **(D)** Representative proteome profiler array membrane images for HUVECs with or without treatment of model CAF sEVs. Red boxes indicate reference spots in duplicate, green boxes indicate negative control spots. **(E)** Heat map of angiogenesis-related protein abundance quantified via integrated density of model CAF sEV treated HUVECs, normalised to untreated. **(F)** Protein abundance of angiogenesis-related proteins of model CAF sEV treated HUVECs, normalised to untreated HUVEC values. Statistical significance was calculated using a one sample Wilcoxon signed-rank test. Quantification of **(G)** α-SMA, **(H)** vimentin, **(I)** SNAIL, **(J)** VE-Cadherin and **(K)** CD31 transcript expression via qPCR in HUVEC cells. Significance was quantified with a one-way ANOVA. Range bars indicate SEM (*, p≤0.05, **, p≤0.01, ***, p≤0.001, ****, p≤0.0001).

### Model CAF sEVs drive re-education in a model of the brain PMN

Melanoma exhibits tropism towards the brain, with around half of all deaths from the disease presenting brain metastases [45]. Various phenotypic and transcriptional events have been previously associated with melanoma brain metastasis, including destabilisation of tight junctions, EndMT and activation of TGFβ/PDGFβ/PDGFRβ signalling [46, 47]. Having shown that co-culture with model CAF sEVs impact HUVECs, we were keen to investigate how they impacted an *in vitro* model of early BBB disruption – a key event in establishment of the brain PMN - using the human brain microvascular endothelial cell line, hCMEC/D3.

Evaluation of pro-angiogenic potential via the tube formation assay revealed that model CAF sEVs enhanced tube formation capacity in hCMEC/D3 cells, with significantly increased branching index and total vessel length per image (Fig. 5A-C + Supplementary Fig. 5A-D). Furthermore, sEVs from both model CAFs and invasive melanoma line A2058 caused a significant increase in both the migratory and invasive properties of the endothelial cells, a phenomenon not observed with NDF sEV treatment (Fig. 5D-E + Supplementary Fig. 5E-F). In addition, mesenchymal EndMT markers were evaluated via qRT-PCR, which revealed increases in model CAF sEV treated samples compared to the untreated control (Fig. 5F-G + Supplementary Fig. 5G-M).

**Figure 5:**
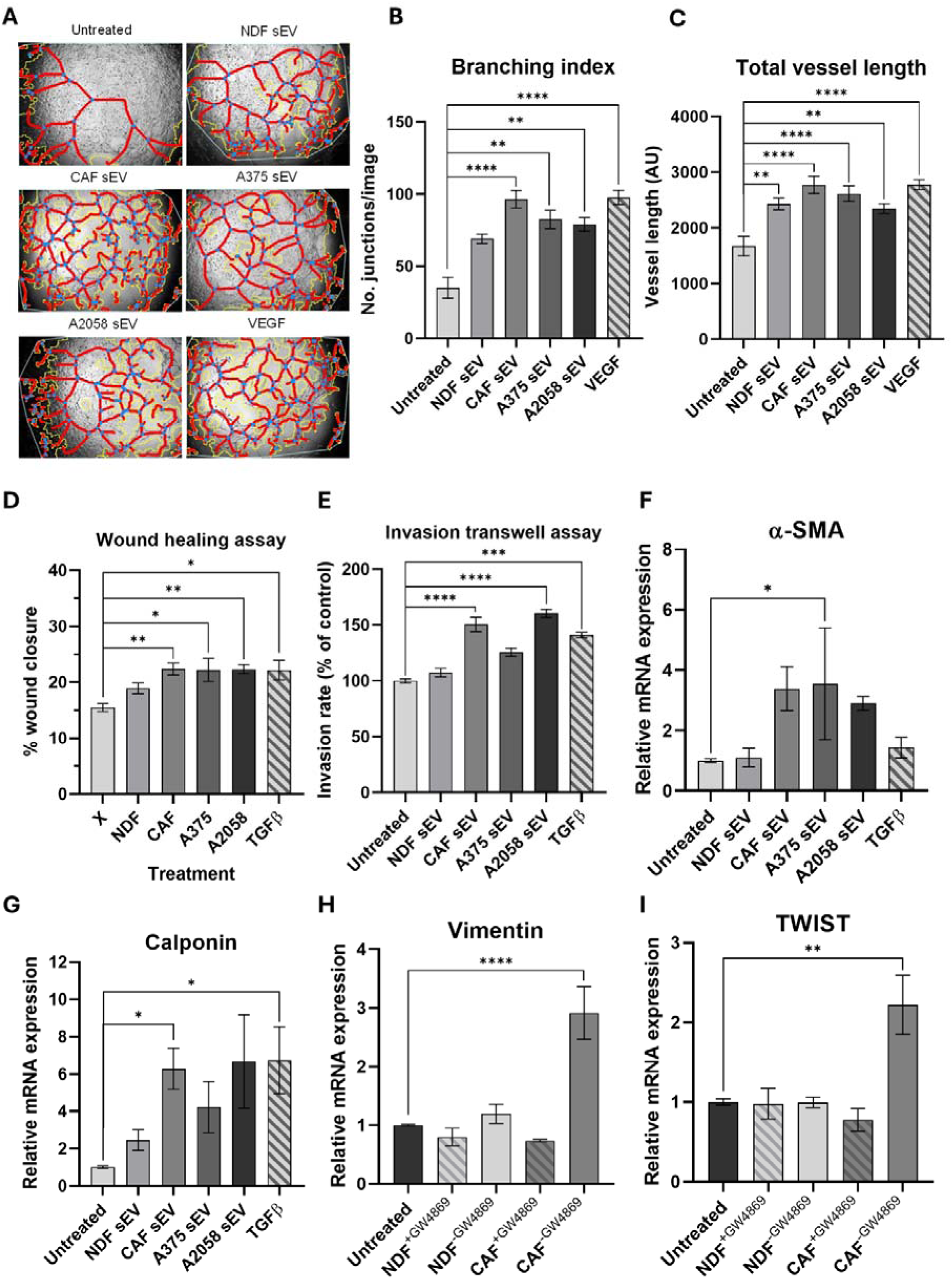
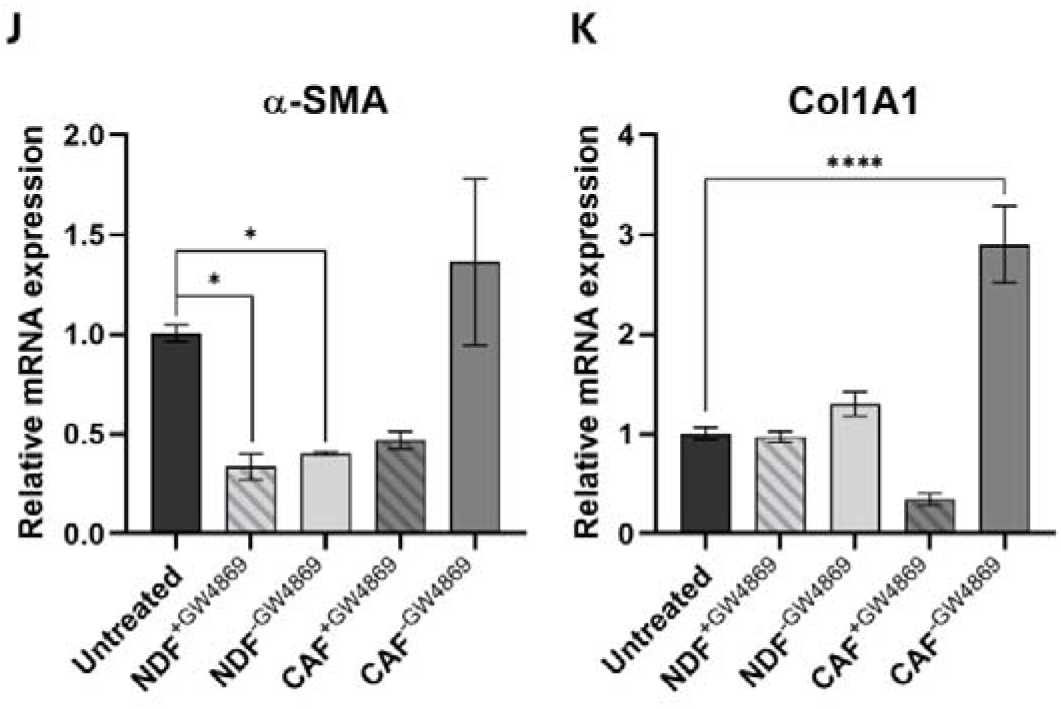
Model CAF sEVs drive re-modelling of brain endothelial cells. **(A)** Representative angiogenesis branching assay images for hCMEC/D3s with treatment of sEVs. Images were captured on the EVOS XL Core Cell Imaging System, at 10x objective and analysed using AngioTool (white line = explant area, yellow line = boundary of vessel, red line = vessel, blue dot = branching point). Quantified **(B)** branching index (no. of branching junctions per image), **(C)** Total vessel length (sum of all vessel lengths in pixels). **(D)** Quantified cell migration as % wound closure in treated hCMEC/D3 cells. **(E)** Quantified cell invasion as % area invading cells normalised to the untreated control. Quantification of **(F)** α-SMA and **(G)** Calponin gene expression via qRT-PCR in hCMEC/D3 cells. Quantification of **(H)** Vimentin, **(I)** TWIST, **(J)** α-SMA and **(K)** Col1A1 gene expression via qRT-PCR in hCMEC/D3 cells cocultured with NDF or model CAFs pre-treated with or without GW4869.

To better understand how continuous exposure to sEVs released by NDFs and CAF model impacted hCMEC/D3 cells, fibroblasts and endothelial cells were co-cultured in a transwell assay, with and without attenuation of sEV secretion. First, we confirmed that the neutral sphingomyelinase inhibitor, GW4869, could successfully attenuate sEV secretion in both NDFs and model CAFs (Supplementary Fig. 5N). We then indirectly co-cultured fibroblasts with and without GW4869 pretreatment with hCMEC/D3 cells and analysed the expression of EndMT markers via qPCR. As can be seen in Fig. 5H-K, we observed a significant increase in vimentin, TWIST and Col1A1 transcripts in hCMEC/D3 cells co-incubated with model CAFs, which was absent in GW4869-treated samples. Together, these data demonstrate that sEV release from model CAFs drive transcriptional and phenotypic changes in an *in vitro* model of the BBB.

### Model CAF-derived small extracellular vesicles are enriched in pro-metastatic cargo and mirror features of melanoma patient plasma

Small extracellular vesicles (sEVs) mediate their functional effects on target cells through several complementary mechanisms, the most widely studies being direct delivery of bioactive cargo, such as mRNAs, non-coding RNA (ncRNAs), proteins, and lipids, which can reprogram gene expression and modulate signalling pathways in recipient cells [48, 49]. To investigate the biomolecular cargo of CAF-derived sEVs and determine how it differs from that of NDF-derived sEVs, we used a multi-omics approach combining quantitative TMT-based LC-MS and small RNA sequencing to assess protein and miRNAs, respectively. TMT LC-MS was performed on sEVs isolated from NDF and CAFs. Following filtering for common contaminants and low-confidence hits, alignment was performed with the EXOCARTA proteome database [50], which confirmed significant enrichment of validated sEV proteins within our dataset (Fig. 6A). Next, we compared normalised abundancies for each sample and identified 19 proteins that were significantly different between NDF and CAF-derived sEVs, with Thrombospondin-1 exhibiting the greatest enrichment in CAF-derived sEVs (Fig. 6B). To explore the functional implications of proteins enriched in CAF-derived sEVs, gene ontology (GO) analysis was performed on EXOCARTA validated proteins, which identified GOs consistent with CAF biology (Supplementary Fig. 6A). Next, we carried out GO analysis for proteins significantly increased in our model CAF-derived sEVs, again we observed GOs associated with CAFs but strikingly GOs associated with vascular regulation were also significantly enriched (Fig. 6C). Next, we performed small RNA sequencing on total RNA from NDF- and CAF-derived sEVs to identify transcripts with significantly different expression levels between the two groups. As can be seen in Fig. 6D, miR-223-3p, miR-148a-3p and miR-34c-5p were enriched and miR-223-5p, miR-1-3p, miR-543 and miR-145-5p were decreased in CAF-derived sEVs compared with NDF control, respectively. Interrogation of our small RNA-seq data also revealed significant differences in other, non-miRNA, genes (Fig 6E), including enrichment of ZC3HAV1. An emerging hypothesis is that sEV released by CAFs (and other tumour cells) may influence the PMN by entering the circulation and impacting on distal tissues [51–53]. To this end, we were interested in determining if any of the protein or genes enriched in our model CAF-derived sEVs were increased in the blood plasma of melanoma patients and thus might have clinical relevance. Melanoma datasets present in The Cancer Genome Atlas (TCGA) and The Human Protein Atlas were of limited utility as these mostly focus on tumour tissue, rather than patient plasma. Instead, we turned to a recent study that reported proteomic profiling of tumour and non-tumour-derived circulating sEVs in the plasma of melanoma patients with patient follow up for progression of disease [54]. Interestingly, this study identified Thrombospondin-1 as significantly enriched in melanoma tumour-derived sEVs, which aligns with our quantitative proteomics analysis of our model CAF-derived sEVs. More strikingly, of the 150-proteins identified by the authors at melanoma-specific Thrombospondin-1 was in the top 5 of a subset of proteins that were predictive for metastatic progression of disease [54]. Next, to ascertain if CAF sEV-enriched RNAs were reflected in clinical datasets, we returned to our previously published transcriptomic data from the primary melanomas of 703 patients, which comprises part of the Leeds Melanoma Cohort (LMC) [55]. Unfortunately, the HT12.4 array utilised in this study does not contain probes for all the ncRNAs identified in our RNA-seq data, but we did observe a significant correlation between expression of ZC3HAV1 and miR-148a and patient survival in primary melanoma tumour samples (Fig. 6F). Interestingly, while increased levels of miR-148a predicted reduced survival, ZC3HAV1 expression exhibited an inverse correlation. As RNA-seq data from melanoma patient plasma is mostly unavailable from TCGA, we revisited a recent study that reported blood plasma miRNA expression data from individuals with invasive melanoma or related benign phenotypes [56] and interrogated these data to investigate if any of the differentially enriched miRNAs identified in our CAF-derived sEVs were significantly altered in melanoma patients with metastatic disease. As can be seen in Fig. 6G, 5/7 miRNAs identified in our RNA-seq data were significantly altered in melanoma patient plasma, with miR-145-5p and miR-34c-5p displaying correlative enrichment in both datasets. Collectively, these data report model CAF-derived sEV-enriched protein and RNA, reveal biological functions for these related to PMN remodelling, and include proteins and genes associated with clinical melanoma studies, and patient outcomes.

**Figure 6:**
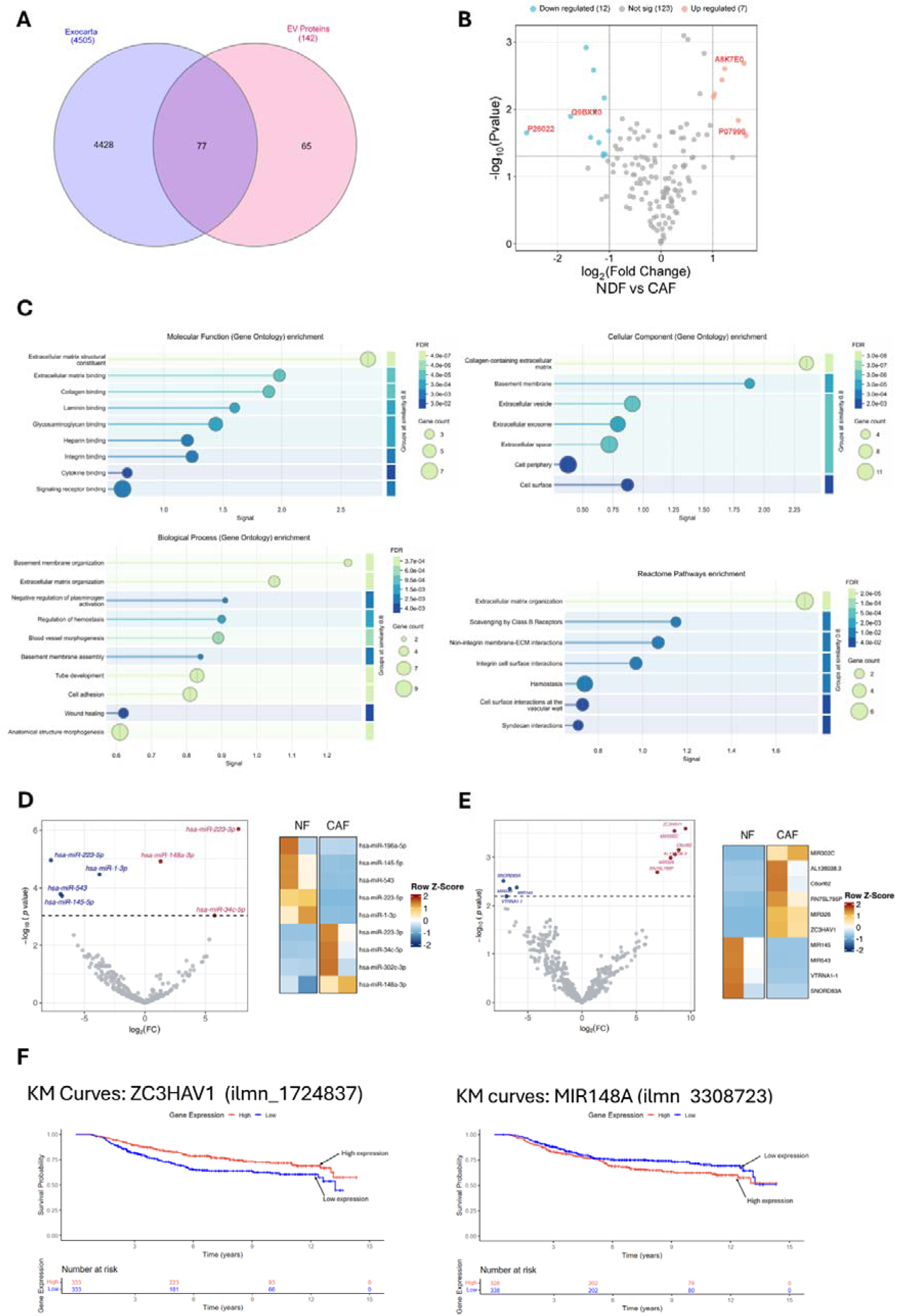
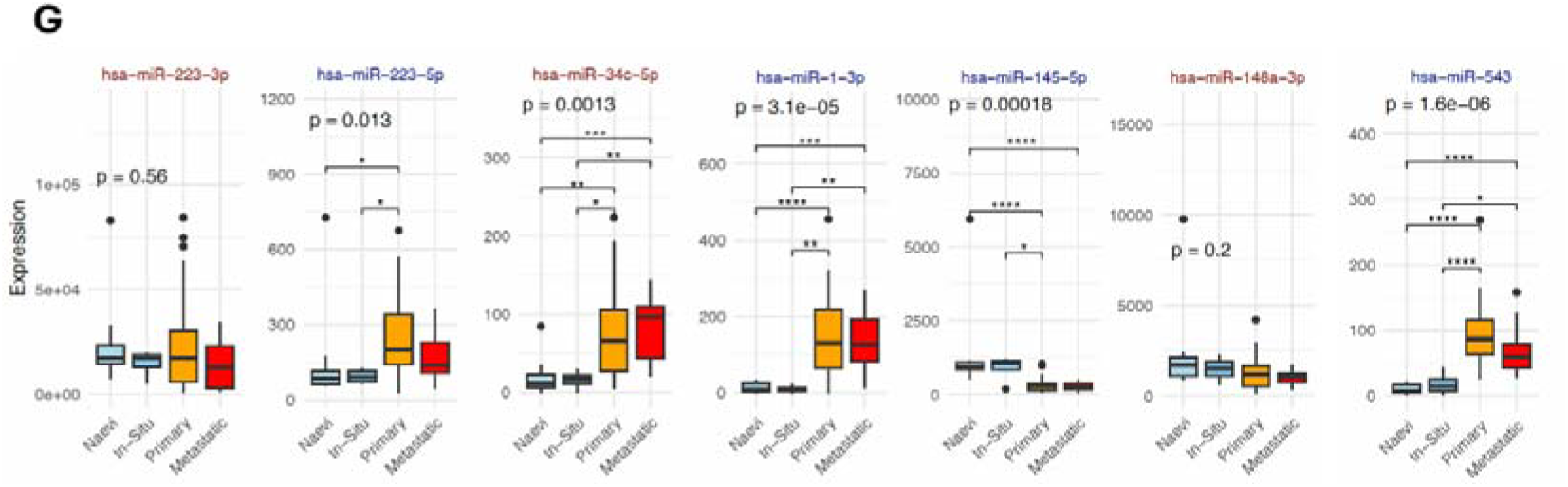
CAF-derived small extracellular vesicles are enriched in pro-metastatic cargo and recapitulate features of melanoma patient plasma-derived sEVs. **(A)** The TMT-labelled proteomic dataset was filtered for high-confidence UniProt-reviewed proteins and compared with the EXOCARTA database, revealing that approximately 54% of the identified proteins were associated with extracellular vesicles. **(B)** Volcano plot of TMT LC-MS analysis of protein abundance from model CAF sEVs compared to NDF sEVs. **(C)** TMT-MS-identified proteins enriched in CAF-derived sEVs were subjected to Gene Ontology (GO) overrepresentation analysis (Biological Process, Molecular Function, and Cellular Component) and Reactome pathway analysis using the PANTHER classification system Enrichment was assessed against the Homo sapiens reference using Fisher’s Exact test with FDR correction. **(D)** Volcano plot of small RNA-seq analysis of miRNA upregulated (red) or downregulated (blue) in model CAF sEVs compared to NDF sEVs (left panel). Z-score hierarchical clustering heat map visualisation of differentially expressed miRNA in model CAF or NDF sEVs (right panel). **(E)** Volcano plot of non-miRNA gene identified in our small RNA-seq, analysed as described above. **(F)** Prognostic value of the top hits from *in-vitro* studies were assessed against the LMC. Kaplan-Meier curves were plotted and Cox proportional hazards regression applied setting high expression as reference. Melanoma-specific survival was analysed ignoring the minority of deaths that were from non-melanoma causes. n=666. **(G)** Previously published data for miRNA-seq analysis of melanoma patient blood plasma (GSE150956) was interrogated to determine transcript levels of miRNA enriched in model CAF-derived sEVs.

## Discussion

CAFs are increasingly recognised as a heterogeneous population whose functions are highly context-dependent, shaped by tumour-intrinsic signals, tissue type, and local environmental cues [36]. In melanoma, where changes to the ECM, immune modulation, and vascular remodelling are fundamental to metastatic spread, CAFs are uniquely poised to support these transitions [57, 58]. Here we opted to investigate how CAF-derived sEVs might contribute to modulation of the TME by establishing a simple, *in vitro* melanoma CAF-like cell model derived from primary dermal fibroblasts and based on previously reported studies [31, 32, 59]. This approach enabled us to work with consistent cell populations, which possessed key phenotypic and marker profiles characterised in the majority of CAFs, as well as successfully isolating and characterising sEVs for downstream analysis and functional assays (Figs. 1 and 2). TGFβ1 was chosen as is a well-established inducer of the myoCAF phenotype [60], however, it represents only one axis of CAF heterogeneity. While our data demonstrate a convincing reprogramming of NDFs, it is important to consider that CAFs derived from alternative cues, for example IL-1 or IL-6 signalling, will likely release sEVs with distinct cargo and therefore comparing sEVs from fibroblasts activated by different tumour signals warrants further investigation.

Whereas early studies on sEV-mediated tumourigenesis focused on unidirectional effects of cancer cell-derived vesicles, contemporary models propose that cells within the TME ‘co-evolve’ via multi-directional crosstalk [61–63]. Our *in vitro* cell line data support this model, specifically the observation that sEV released by our model CAFs not only promote significant increases in mesenchymal-associated phenotypes in melanoma cells but also remodel endothelial cells and non-activated NDFs (Fig. 3 and 4). Importantly, CAF-derived sEVs were not unique in this effect, as widely reported elsewhere melanoma-derived sEVs were also able to drive similar phenotypic changes [64–68]. Interestingly, sEVs isolated from A2058 cells, which are considered to represent a more metastatic model [42, 43], elicited stronger effects compared to those from the less invasive A375 cells suggesting that melanoma cell-derived sEVs vary in their capacity to induce phenotypic changes within the tumour microenvironment. This mutual reinforcement suggests a dynamic co-evolution, potentially establishing a self-perpetuating feedback loop. It highlights the TME as a Gordian knot of intercellular crosstalk, intricately involving cancer, stromal, endothelial and immune cell populations alike [69]. Indeed, longitudinal profiling of co-cultures that incorporate macrophages [70] and other residents immune cells alongside the continued development of microfluid organotypic models [71, 72] will be crucial for fully exploring and elucidating the biology underpinning TME co-evolution and the contribution of sEVs to this process.

Studies investigating the impact of CAF-derived sEVs on endothelial cells are somewhat limited, but not without precedent [73, 74]. We observed a significant activation of angiogenesis in HUVEC cells cultured with CAF-derived sEVs. This effect was similar in magnitude to the responses elicited by A2058 sEVs and VEGF and led to a general increase in pro-angiogenic markers (Fig. 4F). We were particularly interested in how CAF-derived sEVs might impact sites of distal PMN, particularly the brain, and so expanded our analysis to include the human brain microvascular cell line, hCMEC/D3, as a simple model of the blood-brain barrier [75]. We observed similar activation of angiogenesis in hCMEC/D3 cells alongside increases in cell migration and changes in the expression of genes associated with EndMT (Fig. 5). Seminal research demonstrating the tropism of cancer-derived small extracellular vesicles (sEVs) for the lung, liver, and brain [76] laid the groundwork for understanding the contribution of circulating tumour-derived sEVs to the modulation of the PMN. While this concept is now well-established, most subsequent studies have focused on the impact of cancer-derived sEVs [77], including a recent article that describes a role for sEVs released by the brain-seeking variant of triple-negative breast cancer MDA-MB-231 cells in the brain PMN [78]. However, there are relatively few studies considering the role of CAF-derived sEVs in distal PMN remodelling [79], despite the recognition now given to the importance of these cells in disease progression [58, 80–82]. To characterise the protein and miRNA content of CAF-derived sEVs we performed quantitative proteomics and small RNA-sequencing. As expected, our proteomics data was enriched with widely reported sEV proteins, and both NDF and CAFs exhibited GOs associated with ECM organisation, wound healing and cell adhesion. Interestingly, when we investigated GOs based on those proteins that were significantly increased in our model CAFs, several ontologies relating to vascular remodelling reached significance (Fig. 6C). STRING analysis of CAF-enriched proteins revealed a tightly interconnected core of ECM proteins along with modules anchored in vascular remodelling and haemostasis, suggesting that CAF-enriched sEV proteins may fulfil specific functions. Indeed, the protein most significantly enriched in our model CAFs was Thrombospondin-1 (Fig. 6B), which plays dual roles in modulating cell-matrix interaction and angiogenesis with a second angiogenic regulator, PXDN (Vascular peroxidase I), also significantly enriched in our CAF-derived sEVs. The functional roles of these proteins in angiogenesis are tissue-dependent but strikingly both have recently been reported in genome-wide screens to identify genes that significantly correlate with melanoma metastasis [83–86]. Other proteins identified in our TMT-MS dataset that are differentially abundant in CAF-derived sEV include the proteoglycans Decorin and Biglycan, which are depleted and enriched, respectively. Strikingly, there is a growing body of evidence from mouse studies and melanoma patient data that increased Biglycan levels promotes melanoma metastasis [87], drives tumour angiogenesis [88, 89] and correlate with poor patient outcomes [87, 89]. In contrast, Decorin is reported to function in a tumour suppressive manner and low levels in both melanoma tissue and circulating blood plasma have been correlated with disease progression and decreased survival rates in melanoma patients [83, 90].

Analysis of our small RNA-seq data revealed several RNAs that were present at significantly different levels in CAF-derived sEVs compared with NDF controls (Fig. 6C and D). To explore if circulating levels of these genes were altered in melanoma patients we turned to a recent study that published small RNA-sequencing data for 64 plasma biopsy samples from individuals with invasive melanoma or related control [56]. Interrogation of these data for the 7 miRNAs substantively different in our CAF-derived sEV revealed that 5/7 miRNAs exhibited significant differences in abundance in circulating plasma. The direction of expression for these miRNAs was not consistent with changes observed in our CAF-derived sEVs but do hint at a possible panel of miRNA that might warrant further investigation in terms of their diagnostic power. A similar scenario emerged for the non-miRNA genes identified in our small RNA-sequencing data. Here identified genes were not present in the study above and so we leveraged our LMC cohort, which is comprised of primary tumours (35% stage I, 50% stage II and 15% stage III) rather than blood plasma and cause of death and subsequently analysed survival was exclusive to melanoma-associated deaths [55]. Most genes in Fig. 6E were not significant in terms of Hazard Ratios (HR), however, high expression of ZC3HAV1 and MIR148A were inversely and positively correlated with patient survival, respectively. While it is possible to extrapolate GOs from miRNA lists, these data are somewhat problematic as they are generally derived from predicted targets, do not consider how hierarchal interactions might impact on RNA-targets and subsequently their functions and often miss cell-type specific effects. Considering those miRNA significantly increased in CAF-derived sEVs, it is intriguing to note that miR-223-3p, miR-148a-3p, and miR34c-5p, have been reported to be enriched in sEVs derived from a range of different cancers [91–93] and are predominantly reported to function as tumour-suppressors [94–101]. We were somewhat surprised that miRNA increased in CAF-derived sEVs were reported to be tumour-suppressive, however, an emerging concept regarding the release of sEVs within the TME is that cancer cells and CAFs may exploit the packaging of miRNAs into sEVs and their subsequent release to deplete miRNAs that negatively regulate cell growth, thereby promoting a cancer-associated phenotype [102, 103]. Evidence to support this model is on the increase, for example a recent study showed that non-small cell lung cancer cells escape the suppressive effects of the tumour-suppressive miRNA, miR-4732-3p by selectively packaging them into sEVs, thereby supporting tumour progression [104]. How cells selectively export miRNA is still not fully understood, but RNA-binding proteins are likely culprits, hnRNPK implicated in the previous example and YBX1 recently reported to selectively package miR-233 into sEVs [105].

In summary, more work is needed to determine the contribution of CAF-derived sEV to circulating levels of key modulators of the ECM and tumour vascular network. With the development of emerging pipelines for deconvolution of tissue- and cell-type-specific extracellular vesicle abundances [106, 107] we are approaching a point where we can identify which tumour cells contribute significant levels of pro-tumourogenic biomolecules to the circulation. Our data suggest that CAF-derived sEVs may play a significant role in this process, positioning them as an attractive therapeutic target. Through omics analysis of model CAF-derived EVs, combined with phenotypic data on their impact on brain endothelial biology, our data suggest some compelling insights into melanoma brain tropism. It will be crucial to explore how these findings translate to a clinical setting, and we are currently performing longitudinal plasma sampling in melanoma patients to assess whether CAF-specific miRNAs or proteins can serve as predictors of metastatic progression.

## Acknowledgements

The authors would like to thank Dr K. Heesom for assistance with TMT-MS. The Leeds Melanoma Cohort was funded by CRUK grants C588/A19167, C8216/A6129, C588/A10721 and NIH grant CA83115. We thank all the study participants. This work was funded by a British Skin Foundation award to JRB (030/s/18).

