## Supplementary information for "Extracellular vesicles from a model of melanoma cancer-associated fibroblasts induce changes in brain microvascular cells consistent with pre-metastatic niche priming"

**Reviewer access details for TMT-MS data**

Log in to the PRIDE website using the following details:

 Project accession: PXD063635

 Token: sO35VcbiFsjZ

Alternatively, reviewer can access the dataset by logging in to the PRIDE website using the following account details:

 Username: 

 Password: ZNu7ZvyunbTA

**Reviewer access details for RNA-seq data**

RNA-seq data has been deposited with GEO and can be accessed at the location below using the indicated reviewer token.

<https://www.ncbi.nlm.nih.gov/geo/query/acc.cgi?acc=GSE296799>

reviewer token: mlgneqemxrstfkh

**Supplementary methods**

**TEM**

TEM imaging was carried out by University of Leeds Faculty of Biological Sciences, using the FEI Tecnai TF20 microscope: FEGTEM Field emission gun TEM/STEM fitted with HAADF detector, Oxford Instruments INCA 350 EDX system/80mm X-Max SDD detector and Gatan Orius SC600A CCD camera. 5µL drops of sEVs were added to glow discharged TEM grids for 30 sec, blotted with filter paper and washed immediately with drops of EV-free grade 1 filtered water. Samples were stained multiple times with drops of 2% uranyl acetate, blotting between each stain. A final 5µL droplet of uranyl acetate was applied to the grid for 3 min, then blotted to leave a thin film of stain behind. The grid was dried under a lamp for 10 min before imaging.

**RNA extraction**

Cellular RNA extraction was carried out using the Direct-zol™ RNA MicroPrep kit (Zymo Research) according to the manufacturer’s protocol. The isolated RNA 260/280 purity and concentration in ng/µL was quantified using the NanoDrop™ 2000 Spectrophotometer (ThermoFisher). RNA was isolated from 625μL sEVs pretreated with 1mg/mL Proteinase K for 10 min at 56°C, followed by treatment with 4μg/mL of RNase A for 1h at 37°C. Samples were processed using the mirVana™ PARIS™ RNA and Native Protein Purification Kit according to the manufacturer’s protocol for Total RNA (ThermoFisher). 50µL total RNA isolated from sEVs was cleaned up using the RNA Clean & Concentrator™ kit (Zymo) according to manufacturer’s protocol, where small RNA (17-200nt) was further concentrated for RNA sequencing.

**qRT-PCR**

cDNA was synthesised from 300ng-1µg RNA using the iScript™ gDNA Clear cDNA Synthesis Kit (BioRad). cDNA was amplified by RT-PCR using SsoAdvanced™ Universal SYBR^®^ Green Supermix (Biorad). Samples were run in triplicate and analysed using the CFX96™ Real-Time PCR Detection System (BioRad). Transcript expression was determined by analysing a minimum of three biological repeats via geometric averaging [1] of TBP1, GAPDH, and RER1 reference gene to normalise target gene Cq values, providing fold change in gene expression. Primer sequences are found in Table S2.

**Proteome Profiler Human Angiogenesis array**

Expression profiles of angiogenesis-related proteins were carried out on HUVECs treated with or without CAF sEVs for 48h using the Proteome Profiler™ Human Angiogenesis Array (R&D Systems) according to manufacturer’s protocol. 125μg lysate was added to each nitrocellulose membrane in duplicate, and developed on the ChemiDoc™ MP Imaging System (BioRad). Spots were quantified on ImageJ via integrated density after background signal was subtracted, normalised to the reference spot densities.

**Amnis^®^ FlowSight^®^**

sEVs were quantified via NTA and used neat, diluted 1:10 or 1:100 in EV-free PBS (Gibco). sEVs were stained with FITC Anti-CD63 antibody [MEM-259] (Abcam) at a 1:20 dilution that was predetermined as optimal, with appropriate controls (sEVs without FITC Anti-CD63 and EV-free PBS with or without FITC Anti-CD63) according to MIFlowCyt-EV guidelines for standardised reporting of EV flow cytometry experiments. 200-500μL samples were run on the Amnis^®^ FlowSight^®^ Flow Cytometer (Luminex) with a 488nm laser for 5 mins at flow rate of 1.2μL/min after QC calibration with FlowSight system calibration reagent. Laser power was adjusted so the Raw Max Pixel Intensity was between the recommended range of 100-4000 counts. Area (pixels in square microns) and Aspect ratio (width to height ratio) was measured and sEVs were gated from larger or non-circular particles on FlowSight software (INSPIRE). Area was plotted against channel intensity for FITC (sum of all pixel intensities with the background subtracted) for the gated particles to determine the amount of CD63-FITC positive sEVs were present in each sample (PBS with FITC Anti-CD63 used to determine background noise).

**Fluorescence polarisation anisotropy**

Fluorescence polarisation was quantified as the difference between the emission fluorescence intensity parallel and perpendicular to the excitation light plane, divided by total emission fluorescence intensity. Fluorescence polarisation was measured using the CLARIOstar Microplate Reader (BMG Labtech) using MARS Data Analysis software (BMG Labtech). 100μL sEVs or EV-free PBS as a control were primed into the reagent injector and 50μL CellMask™ Orange diluted 1:10,000 in EV-free PBS was loaded into wells of a 96-Well Solid Black Polystyrene Microplate (Corning). Gain and focus adjustment settings were carried out between each sample with target millipolarisation (mP) of 35 and gain between 1000-2000, focal height 4.9mm. Well mode settings used were:

-	Kinetic window 1 (0-31.32 sec): 30 intervals, 1.08 sec intervals, 50 flashes per interval

-	Kinetic window 2 (40-71.32 sec): 30 intervals, 1.08 sec intervals, 50 flashes per interval

-	Kinetic window 3 (72.4-190.4 sec): 60 intervals, 2.00 sec intervals, 50 flashes per interval

Optics settings were excitation 540nm and emission 590nm with LP 566 dichroic filter. Injection settings used for sEVs into the CellMask™ Orange were 50μL injection volume at 35 sec at a pump speed of 50μL/sec. Difference in fluorescence polarisation was calculated using the mP values before and after injection.

**Micro-BCA assay**

The Pierce™ BCA Protein Assay Kit (ThermoFisher) was used with sEVs diluted in the range of 1:2-1:20 in 0.1M NaOH. 150µL of each standard or sample was added in triplicate to a microplate well, and 150µL of working reagent (reagents A, B and C at a ratio of 25:24:1) was added and incubated for 2 hours at 37°C. Absorbance was read at 560-570nm using the Tecan Infinite F50 Robotic absorbance plate reader with Magellan Data Analysis software.

**Flow cytometry**

Cells were fixed in ice cold 70% ethanol or 2% paraformaldehyde overnight at 4°C. Cells were washed then resuspended in permeabilization buffer (0.1% Triton™ X-100 (Sigma)) and incubated 10-20 min at room temp. Cells were washed, then resuspended at 1-2 x 10^6^ cells/mL in blocking buffer (0.5% Bovine Serum Albumin (Sigma) + 2% FBS) and incubated at 4°C 30 min. Cells were resuspended in FACs buffer (0.5% BSA + 0.05% NaN_3_ in PBS) with or without a primary antibody at recommended dilution at 4°C 30 mins. Cells were washed in FACs buffer, then resuspended in FACs buffer with or without appropriate secondary antibody at 4°C 30 min, protected from light. Cells were strained to remove doublets using a 70μm pluriStrainer^®^ (PluriSelect). FACs analysis was completed using the BD Accuri™ C6 Flow Cytometer (BD Biosciences). Flow rate was set to low, and 10,000 singlet events were captured after gating for singlets using FSC-A vs FSC-H scatter plot. Data was collected using the appropriate laser for the fluorophore used with standard filter (FL1 = 533nm, FL2 = 585nm, FL3 = 670nm, FL4 = 675nm+). Antibodies used can be found in Table S1

**Immunocytochemistry**

8 well detachable Thermo Scientific™ Nunc™ Lab-Tek™ Chamber Slide System (ThermoFisher) were coated for 1h with 0.01% Poly-L-lysine (Sigma). Slides were washed with PBS and cells were seeded 10^3^-10^5^ per well and left overnight to attach. Samples were fixed in 4% paraformaldehyde for 10 min, rinsed in PBS then incubated with 0.1% Triton™ X-100 (Sigma) for 10 min. Cells were washed then incubated with blocking buffer (1% BSA (Sigma), + 0.1% Tween-20) for 1h. Slides were incubated with primary antibody in PBST-BSA overnight at 4°C. Cells were washed and incubated with secondary antibody if primary antibody was not conjugated to a fluorophore in PBST-BSA for 1h. Cells were washed then mounted in VECTASHIELD^®^ Antifade Mounting Medium with DAPI (Vector Laboratories) after detaching the chamber slide wells. Slides were imaged on the Zeiss Axio Imager M2 (Zeiss) using ZEN Blue 2.0 software (Zeiss). Images were analysed on ImageJ to calculate the corrected total cell fluorescence (CTCF = Integrated Density – (Area of selected cell * Mean fluorescence of background readings)).

**SDS PAGE + Western Blot**

Whole cell lysates and sEV protein extracts were generated by lysis with RIPA Buffer (ThermoFisher) containing Halt™ Protease Inhibitor Cocktail (100X) (ThermoFisher). Lysates were incubated on ice for 30 mins, then centrifuged to remove insoluble material. An equal volume of Laemmli 2x Sample Buffer (Sigma Aldrich) was added and samples were heated to 95°C for 5 mins. For non-reduced samples, Pierce™ LDS Sample Buffer, Non-Reducing (4X) was added (ThermoFisher). 2-15µL samples and 5µL PageRuler™ Prestained protein ladder (Thermo Fisher Scientific) were run on a Mini-PROTEAN^®^ TGX Stain-Free™ Precast Gel (BioRad) and transferred onto Amersham™ Protran^®^ Western blotting nitrocellulose membranes (Sigma Aldrich). Membranes were blocked with 5% TBS-T + milk, then incubated with primary antibodies overnight (see Table S1). Membranes were incubated with secondary antibody then equal volumes of EZ-ECL solutions A and B (Biological Industries) and developed on the ChemiDoc™ MP Imaging System (BioRad).

**Cell morphology image analysis**

Multiple high-power images of cells were taken using the EVOS XL Core Cell Imaging System, at 10x objective. Cell edges were drawn manually using photo editing software and a mask was applied using ImageJ. ImageJ used to measure cell morphology parameters Cell Area (area of selection in square pixels), Cell Circularity (4π(area/perimeter^2^) and Cell Perimeter (length of the outside boundary of the selection).

**Cell migration assay**

Cells were either treated before growing to a confluent monolayer or treated once they reached confluency in serum media. Media was then changed to serum starve cells for 24h prior to the assay. Cells were scraped using a p200 pipette tip to create a straight line and washed to remove detached cells. 0h image was taken on the EVOS XL Core Cell Imaging System, 4x objective. Cells were incubated at 37°C for 6-24h depending on the cell type, and images were taken at routine time points. Images were quantitatively analysed using ImageJ to measure the area between the scratch lines.

**Cell viability assay**

Cells were pelleted and 100μL cell suspension (3-5 x 10^4^ cells per well) were added in triplicate to a white 96 well plate. Plate was incubated for 30 min at room temp, then 100μL CellTiter-Glo^®^ 2.0 reagent was added to each well. Cells were lysed for 2 min on a plate shaker, incubated for 10 min to stabilise luminescent signal, then luminescence was recorded using the CLARIOstar Microplate Reader (BMG Labtech) using MARS Data Analysis software (BMG Labtech). For WST-1 assay, 10μL of Cell Proliferation Reagent WST-1 (Roche) was added per well and mixed thoroughly on a plate shaker. Cells were incubated for 4h, mixed again and absorbance was measured at 450nm using the CLARIOstar Microplate Reader (BMG Labtech) using MARS Data Analysis software (BMG Labtech). A reference wavelength of 630nm was also measured. Actual Abs = (Sample Abs_450_ – Average blank Abs_450_) – (Sample Abs_630_ – Average blank Abs_630_)

**Agarose gel electrophoresis**

0.7-1g agarose (ThermoFisher) was dissolved in 100mL 1x TAE Electrophoresis Buffer (ThermoFisher), ethidium bromide (Sigma) was added to get a concentration of 0.5μg/mL, then poured into a multiSUB^®^ Electrophoresis System (Cleaver Scientific Ltd) to set for 20 min. 1x TAE was added to the agarose gel buffer tank and 10μL RNA samples were run alongside DNA/RNA ladder (Sigma) diluted in DNA/RNA loading dye (ThermoFisher). Gels were run at 100V for 40-60 min then visualised on the ChemiDoc™ MP Imaging System (BioRad).

**Acrylamide gel**

15mL of denaturing 15% polyacrylamide gel with 8M of urea was prepared (7.2g Urea, 1.5mL 10x TBE, 5.6mL 40% acrylamide (acryl:bis acryl = 19:1), nuclease free water, 75µL 10% APS, 15µL TEMED) and poured into Mini-PROTEAN^®^ Tetra Handcast Systems (Biorad). Equal volumes of sEV RNA (total, large or small) were mixed with 2x RNA loading dye (NEB) and heated to 95°C for 5 mins, then immediately put into ice for 3 mins before loading onto gel. 25µL of sample were run, alongside Low Range ssRNA Ladder (NEB) diluted 1:20 in loading dye. Gels were run with 1x TBE at 30-45mA for 40-60 min. Gels were then washed for 60 min in TE buffer pH 8 with SYBR™ Gold Nucleic Acid Gel Stain (Invitrogen) diluted 1:10,000. Images were taken using the ChemiDoc™ MP Imaging System (BioRad) SYBR™ Gold 590/110 application.

**miRCURY LNA miRNA PCR**

cDNA was synthesised with the miRCURY LNA miRNA PCR starter kit (Qiagen) using 2-6μL of RNA per reaction, set up according to the manufacturer’s protocol with UniSp6 spike in added as a reverse transcription positive control. LNA-enhanced primer sets used; miR-21, miR-155-5p, miR-146-5p, miR-221-3p, miR-345-5p, miR-29b-3p, miR-16-5p, and UniSp6 as a control, and were normalised to miR-103a-3p as a reference target.

**Hazard Ratio calculations**

Gene expression profiles (accession number EGAS00001002922) from the Leeds Melanoma Cohort (LMC, ethical approval MREC 1/3/57, PIAG 3–09(d)/2003) were used to test the prognostic value of the top hits from in-vitro studies. The LMC is a large population-based treatment naïve primary melanoma dataset with a long follow up [2, 3]. After splitting gene expressions in high vs. low by median, Kaplan-Meier curves were plotted and Cox proportional hazards regression applied setting high expression as reference. Melanoma-specific survival was analysed ignoring the minority of deaths that were from non-melanoma causes (final sample size=666). Analyses were conducted in R.

**Supplementary tables**

| Antibody | Supplier | Dilution ratio (v/v) |
| --- | --- | --- |
| GAPDH Monoclonal Antibody | Proteintech (60004-1-Ig) | 1:20,000 |
| β-Tubulin Antibody | Cell Signalling Technology (2146) | 1:1000 |
| Recombinant Anti-CD81 Antibody | Abcam (ab109201) | 1:1000 |
| Recombinant Anti-CD63 Antibody | Abcam (ab134045) | 1:1000 |
| Anti-Calnexin Antibody | Abcam (ab22595) | 1:1000 |
| Alpha-Smooth Muscle Actin Monoclonal Antibody | ThermoFisher (MA5-11547) | 1:250 |
| Goat anti-Rabbit IgG (H+L) Secondary Antibody, HRP | ThermoFisher (31460) | 1:10,000 |
| Rabbit anti-Mouse IgG (H+L) Secondary Antibody | ThermoFisher (A16163) | 1:10,000 |
| Goat IgG HRP-conjugated Antibody | R&D Systems (HAF017) | 1:1000 |
| CoraLite488 smooth muscle actin specific rabbit polyAB | Proteintech (CL488-55135) | 1:100 |
| coraLite488-conjugated MMP2 mouse McAb | Proteintech (CL488-66366) | 1:100 |
| Goat Anti-Rabbit IgG H&L (Alexa Fluor® 488) | Abcam (ab150077) | 1:200 |
| Goat Anti-Mouse IgG H&L (Alexa Fluor® 647) | Abcam (ab150115) | 1:200 |

**Supplementary Table S1**. List of antibodies used in this study.

| Oligonucleotide (all Sigma Aldrich) | Sense | Antisense |
| --- | --- | --- |
| GAPDH (ref) | CGGAGTCAACGGATTTGGTC | GGATCTCGCTCCTGGAAGATG |
| TBP (ref) | TTGGGTTTTCCAGCTAAGTTCT | CCAGGAAATAACTCTGGCTCA |
| RER1 (ref) | CGTAGCGGAGCTGCGAG | CGTGTAGGGTGTGGACTTGT |
| HNRNPL (ref) | ACAAACCCCAATCTCAGTGG | CCCTCATCATGGTAATGGCT |
| ACTB (ref) | GGGAAATCGTGCGTGACATT | GACTCCATGCCCAGGAAGG |
| α-SMA | GAAGAAGAGGACAGCACTG | TCCCATTCCCACCATCAC |
| FN1-EDA | TGGAACCCAGTCCACAGCTATT | GTCTTCTCCTTGGGGGTCACC |
| COL1A1 | GTGGCCATCCAGCTGACC | AGTGGTAGGTGATGTTCTGGGAG |
| MMP2 | AATAATTCCGCTTCCAGGGCACATCC | TTATTGCGGTCGTAGTCCTCAGTGGT |
| Vimentin | AACCGACACTCCTACAAG | TTATTGAAGCAGAACCAAGTT |
| TWIST | ACCATCCTCACACCTCTG | GATTGGCACGACCTCTTG |
| SNAIL | GTGTCTCCCAGAACTATT | GTTTGAAATATAAATACCAGTGT |
| VE. Cadherin | CGCAATAGACAAGGACAT | GCCGTGTTATCGTGATTA |
| Calponin | CCAGGCTCCGTGAAGAAG | CTTGATGAAGTTGCCGATGTT |
| CD31 | TCTGATTGGCTAACTGAA | TGAGTGAATGTTGACCTA |
| FSP1 | TACTGTGTCTTCCTGTCCT | TTCCTGGGCTGCTTATCT |

**Supplementary Table S2**. List of oligonucleotides used in this study.

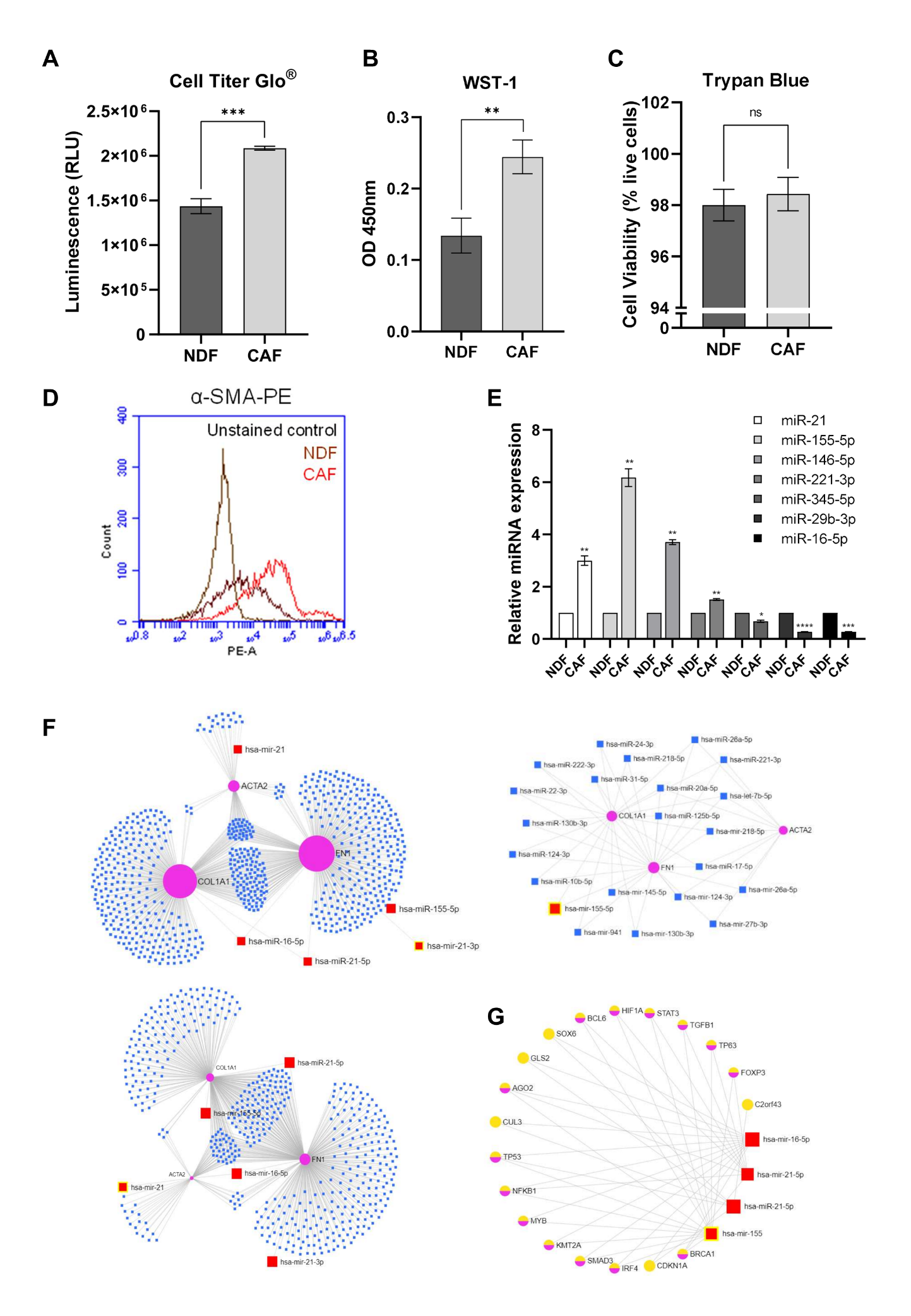
**Supplementary Figure 1: Characterisation of model CAFs treated with TGFβ1. (A)** Quantified relative luminescence units (RLU) of Cell Titer Glo^®^ Luminescent Cell Viability assay in NDFs and model CAFs. **(B)** Corrected optical density at 450nm using 630nm as a reference to measure formazan dye production in NDFs and model CAFs. **(C)** Quantified cell viability as % live cells in NDFs and model CAFs. **(D)** α-SMA-PE of model CAFs vs NDFs compared to unstained controls. Cells were gated around singlets and 10,000 events were collected. **(E)** miRCURY LNA PCR assay quantification for miRNAs commonly up- or down-regulated in CAFs, normalised against miR-103a. **(F)** miRNet miRNA-gene network using 4 CAF-related target genes in all tissue types (upper-left panel), skin tissue (upper-right panel) and exosomes (lower-left panel). Genes with interacting miRNAs and target miRNAs are highlighted in red circles or red squares, respectively. **(G)** miRNet miRNA-gene network of all genes and transcription factors that interact with 3 or more of the target miRNAs (degree filter = 3). miRNAs are highlighted in red squares, genes in pink circles and transcription factors in orange and pink circles.

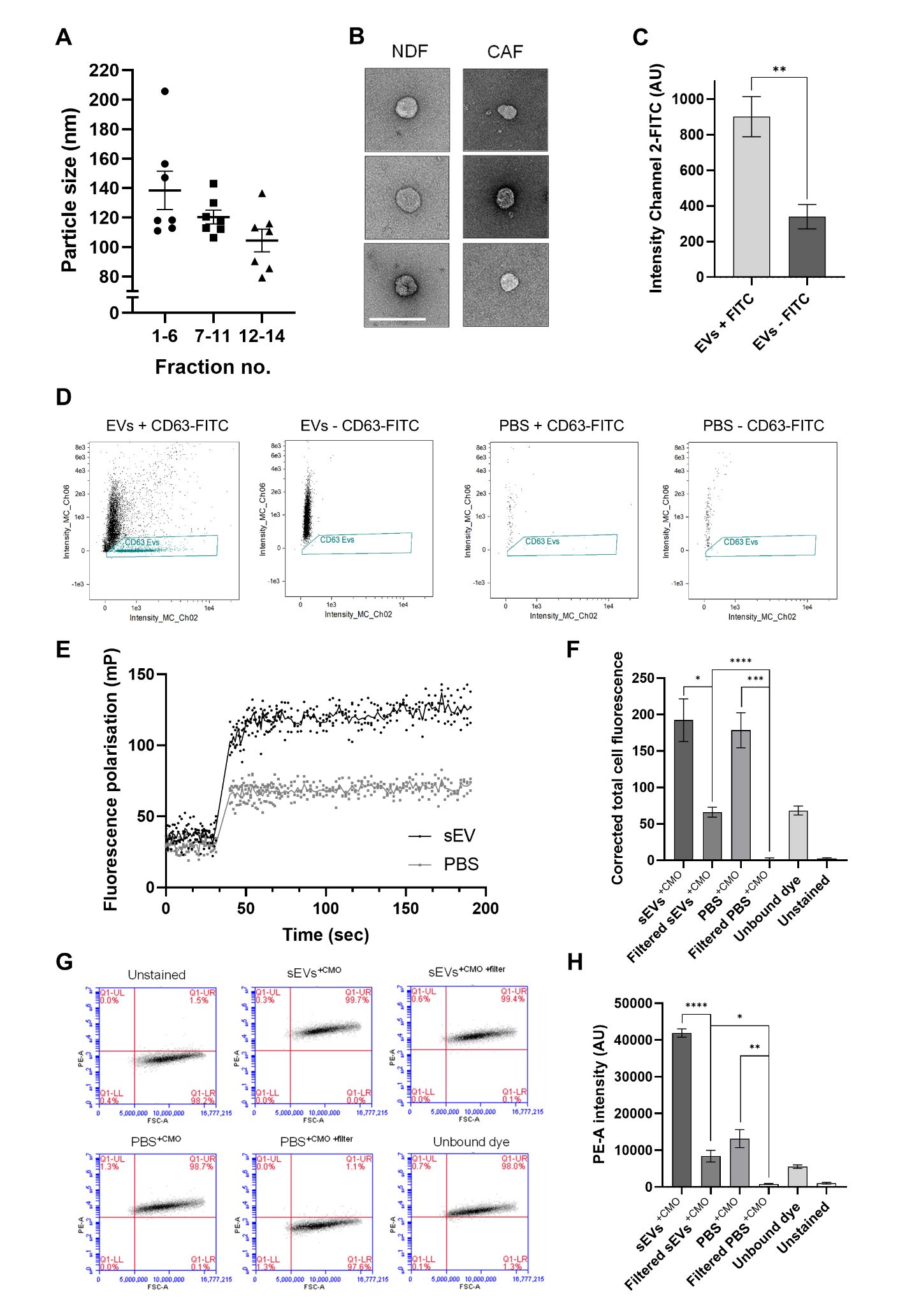

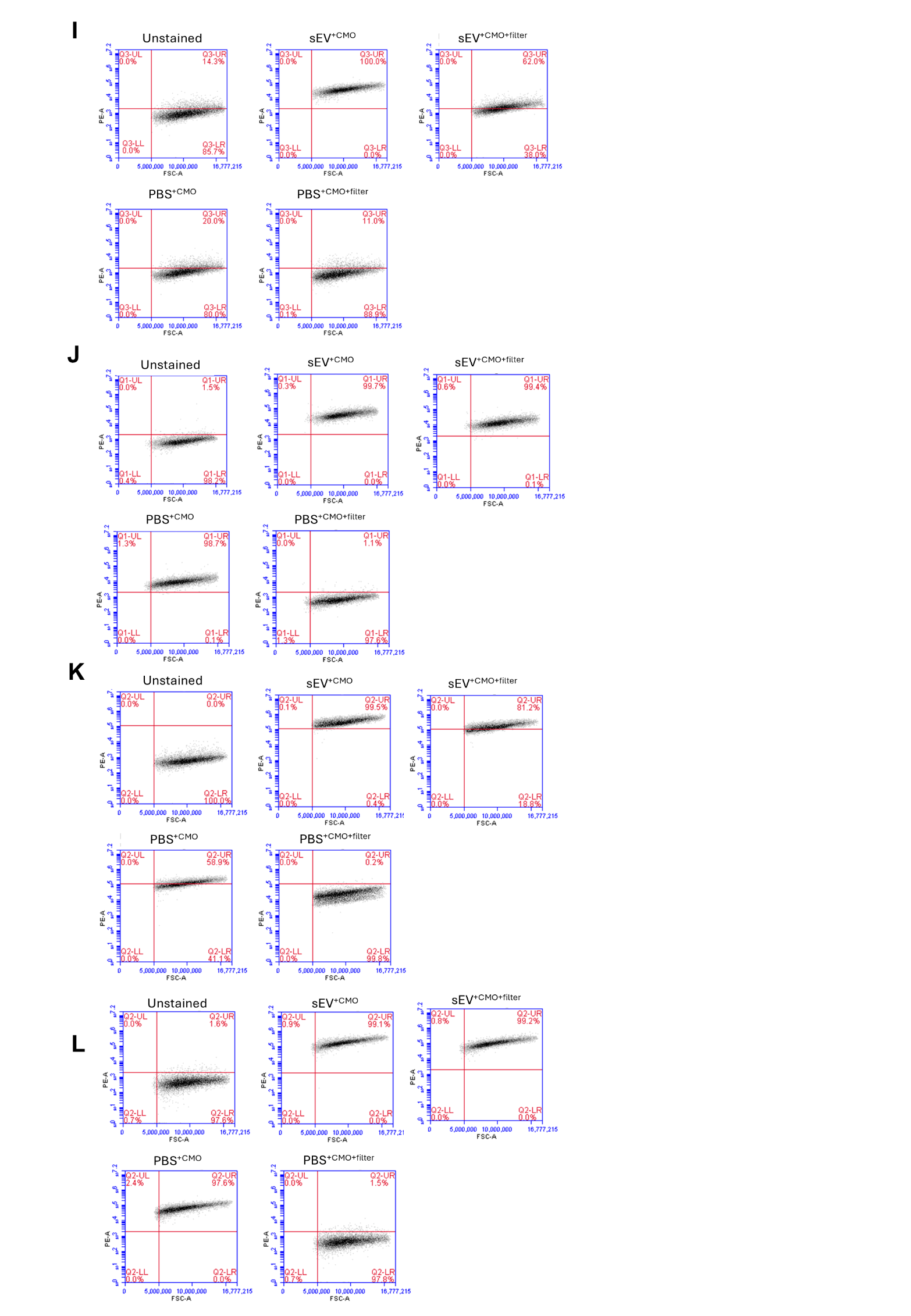
**Supplementary Figure 2: Characterisation of CAF sEVs isolated via SEC and their uptake into recipient cells. (A)** Average particle size in of combined fractions 1-6, 7-11 and 12-14. **(B)** Representative Transmission Electron Microscopy (TEM) images of 5μL of NDF or model CAF isolated combined fractions 7-11 (scale bar = 200nm). **(C)** FlowSight data of sEVs or EV-free PBS stained with or without CD63-FITC after gating for EVs based on area and aspect ratio. Samples were run for a time limit of 5 minutes. **(D)** Average intensity of channel-02 (FITC) in arbitrary units (AU) for sEVs stained with or without CD63-FITC. **(E)** Fluorescence polarisation of CMO dye pre- and post-injection with model CAF sEVs or EV-free PBS. 50µL samples were injected at 35 seconds and polarisation of the emitted light at 590nm was measured in milli polarisation. **(F)** Quantified corrected total cell fluorescence intensity (AU) of IF images of A375 cells treated with model CAF sEVs or PBS with or without CMO, accounting for mean fluorescence of background readings. **(G)** Fluorescence intensity plotted against FSC-A after gating for singlets in A375s treated with sEVs or PBS with or without removal of unbound dye, or unbound dye only. **(H)** Fluorescence intensity for each A375 treatment as PE-A. **(I-L)** Fluorescence intensity plotted against FSC-A after gating for singlets in NDF (I), A2058 (J) , HUVEC (K) and hCMEC/D3 (L) cells treated with sEVs or PBS with or without removal of unbound dye.

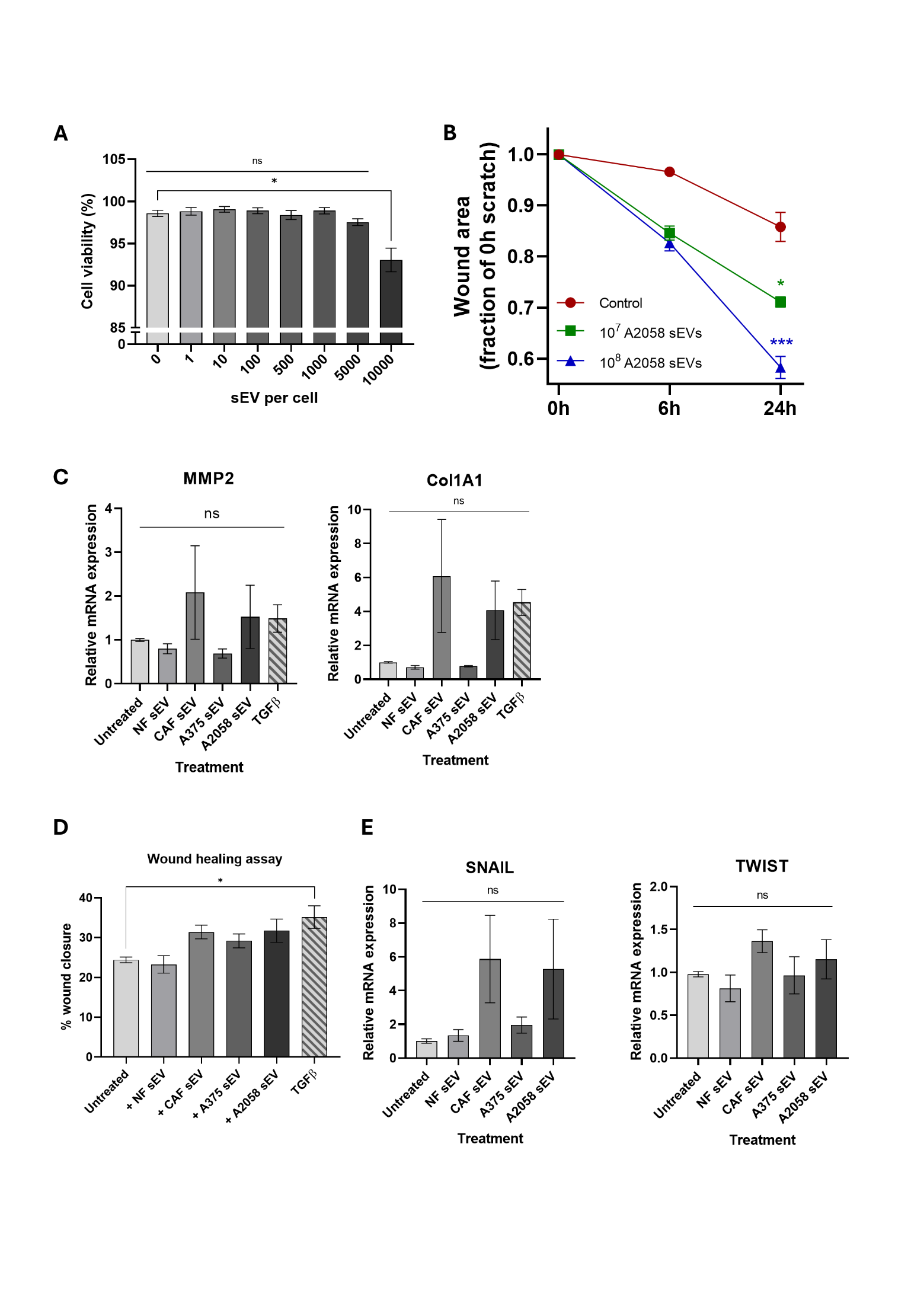
**Supplementary Figure 3: CAF sEV treatment affects multiple phenotypic parameters in dermal fibroblasts and melanoma cell line A375. (A)** Quantified cell viability as % live NDFs post-treatment with indicated number of sEVs. **(B)** Cell migration in NDFs treat with increasing amount of sEVs. **(C)** Relative transcript expression of CAF markers MMP2 and Col1A1 post-treatment with sEVs. **(D)** A375 cell migration as % wound closure at 24h compared with untreated control. **(E)** Relative transcript expression of EMT-associated genes SNAIL and TWIST in A375 cells post-treatment with sEVs.

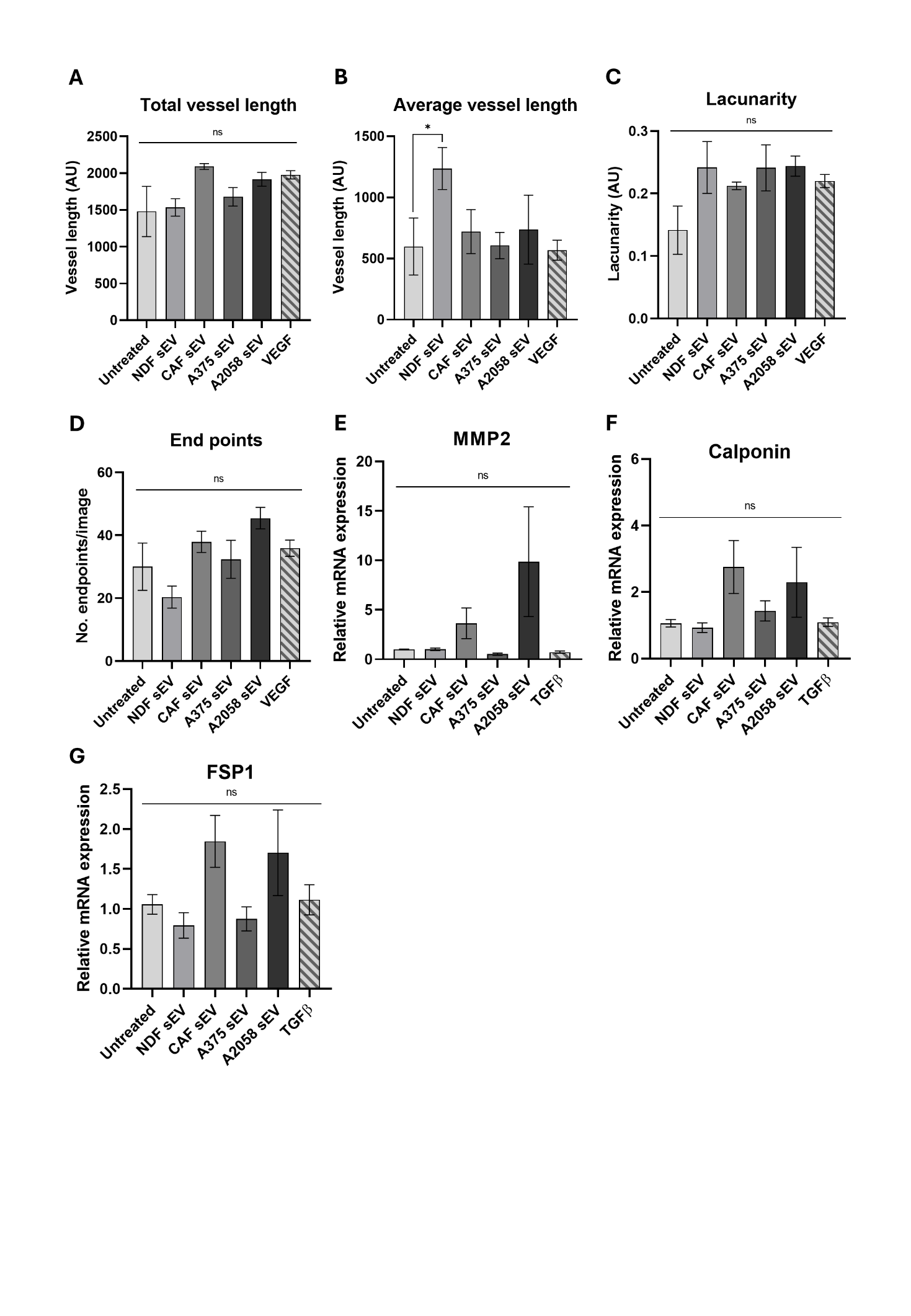
**Supplementary Figure 4: Model CAF sEV treatment changes angiogenic properties of HUVECs.** Quantified **(A)** Total vessel length (sum of all vessel lengths in pixels), **(B)** Average vessel length (mean length of all vessels per image), **(C)** Lacunarity (average lacunarity per image), and **(D)** End points (no. of endpoints per image). Statistical significance was determined for using a one-way ANOVA or Kruskal-Wallis ANOVA. Quantification of **(E)** MMP2, **(F)** calponin and **(G)** FSP1 transcript expression via qPCR in HUVEC cells. Significance was quantified with a one-way ANOVA. Range bars indicate SEM (*, p≤0.05, **, p≤0.01, ***, p≤0.001, ****, p≤0.0001).

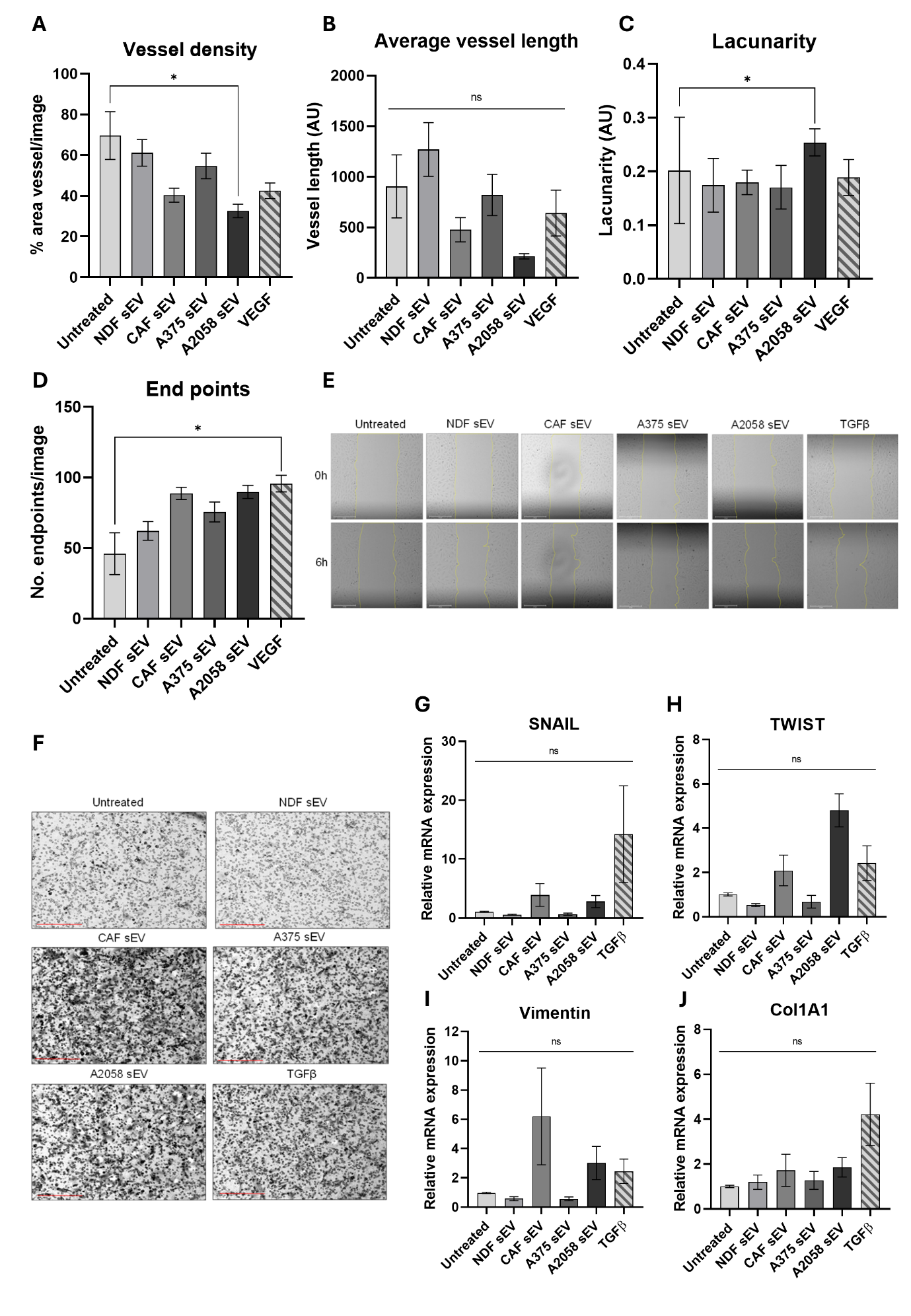

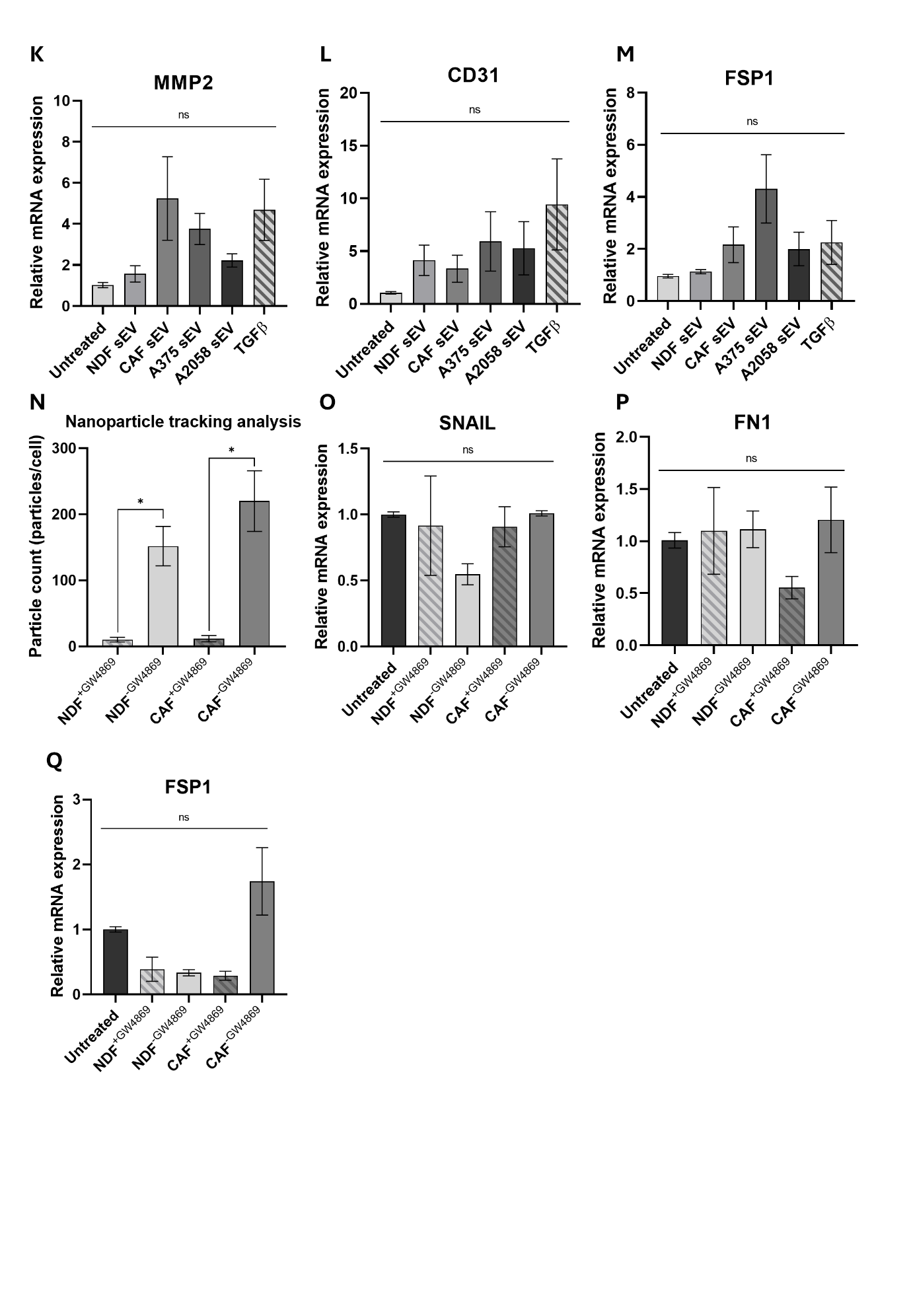
**Supplementary Figure 5: Model CAF sEVs drive re-modelling of brain endothelial cells. (A)** Quantified vessel density (% area in explant area occupied by vessels), **(B)** Average vessel length (mean length of all vessels per image), **(C)** Lacunarity (average lacunarity per image), and **(D)** End points (no. of endpoints per image). **(E)** Representative images of scratch wound healing assay of hCMEC/D3s at the start (0h) and end (6h) of assay (scale bar = 275µm). **(F)** Representative images of a transwell invasion assay of hCMEC/D3s at 24h (scale bar = 275μm). Quantification of **(G)** SNAIL, **(H)** TWIST, **(I)** Vimentin, **(J)** Col1A1, **(K)** MMP2, **(L)** CD31, **(M)** FSP1 gene expression via qPCR in hCMEC/D3 cells. **(N)** Particle count per cell of NDF and model CAFs pre-treated with or without GW4869 quantified via NTA. Quantification of **(O)** SNAIL, **(P)** FN1 and **(Q)** FSP1 gene expression via qRT-PCR in hCMEC/D3 cells cocultured with NDF or model CAFs pre-treated with or without GW4869.

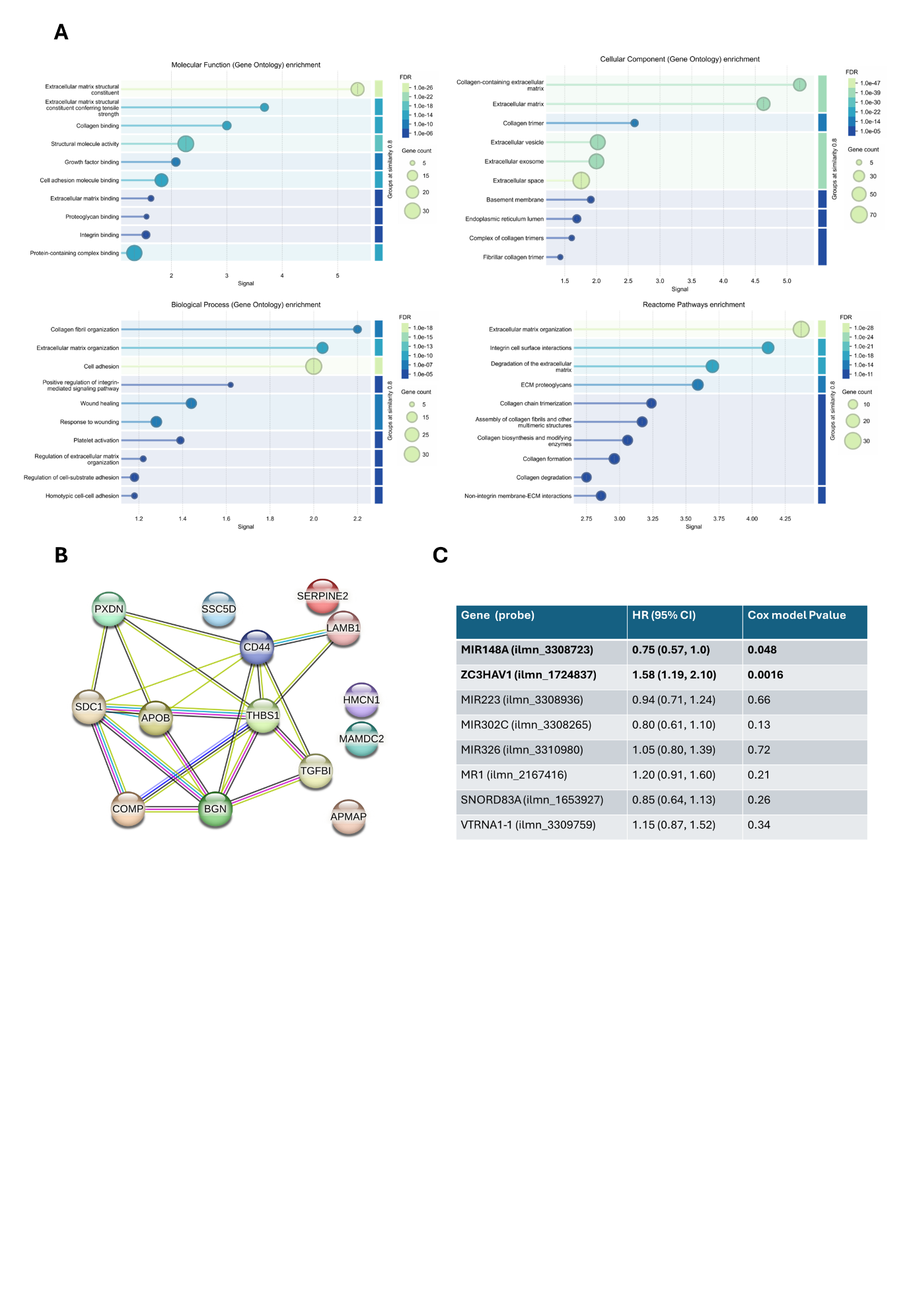
**Supplementary Figure 6: CAF-derived small extracellular vesicles are enriched in pro-metastatic cargo and recapitulate features of melanoma patient plasma-derived sEVs.** **(A)** TMT-MS-identified proteins present in CAF-derived sEVs were subjected to Gene Ontology (GO) overrepresentation analysis (Biological Process, Molecular Function, and Cellular Component) and Reactome pathway analysis using the PANTHER classification system Enrichment was assessed against the Homo sapiens reference using Fisher’s Exact test with FDR correction. **(B)** STRING network analysis proteins enriched in model CAF-derived sEVs [PPI enrichment p-value: 1.33e-06] (red = gene fusion, light purple = experiments, dark purple/black = co-expression, light blue = homology, dark blue = cooccurrence, light green/yellow = text-mining, dark green = neighbourhood, turquoise = databases). **(C)** Cox proportional hazards regression for genes identified in Fig. 6E utilising data from the LMC, n=666.
